# Spatially resolved fetal and maternal cell contributions to severe Preeclampsia

**DOI:** 10.1101/2025.07.18.665528

**Authors:** Yara E Sanchez-Corrales, Theodoros Xenakis, Jose J Moreno-Villena, Leysa Forrest, Neil J Sebire, Elizabeth C Rosser, Lucy R Wedderburn, Sergi Castellano, Sara L Hillman

## Abstract

The molecular and cellular pathophysiology of the fetal-maternal interface in preeclamsia remains poorly understood, but it is increasingly clear that both fetus and mother make independent contributions to it. Here, we spatially distinguish them from each other and from gestational age for the severe form of the disease, interrogating tissues and cell types beyond the placenta for the Early and Late presentation of the condition. In addition to concerted hypoxia, angiogenic imbalance, fibrosis and aberrant metabolism in the placenta, new maternal immune signatures, including changes in macrophages, mitochondrial dysfunction and interferon signalling in the placenta, myometrium or chorioamniotic membranes, offer an explanation to systemic inflammation and endothelial dysfunction in the mother. These tissue and cell-specific responses are potential targets for therapy, with their prompt consideration in Early gestational disease likely beneficial due to its severity. Thus, timely intervention during gestation could change the extremely poor prognosis of severe disease.

## Introduction

The fetal maternal interface (FMI) is the site of interaction between specialised fetal cells (trophoblasts) and maternal cells during pregnancy^1^. It includes the placental villi, where trophoblasts are in contact with maternal blood, and extends to the adjacent decidua basalis (at the site of placentation) and decidua parietalis (surrounding the fetus), where trophoblasts contact maternal cells from the modified endometrium. These interactions remain poorly defined but they are essential to understand, prevent and treat pregnancy complications such as preeclampsia (PE).

PE is recognised as a placental pathology^2^ with systemic consequences, including new onset maternal hypertension and proteinuria at or after 20 weeks of gestation^3^. It is a leading cause of maternal and fetal morbidity and mortality, affecting 2-4% of pregnancies globally with over 46,000 maternal and 500,000 fetal and neonatal deaths per year^4^. Early in pregnancy, the placental cytotrophoblast (CTBs) cells differentiate into syncytiotrophoblasts (STBs), which contact maternal blood, or extravillous trophoblasts (EVTs), which invade through the decidua and remodel the spiral arteries of the myometrium^5^. PE is associated with a shallow EVT invasion and defective spiral artery remodelling leading to placental stress and ischaemia, as well as systemic maternal endothelial dysfunction and endovascular inflammation^3,6^. Importantly, PE is a highly heterogenous disease for which no effective treatment exists^3^.

Recent studies have applied single cell transcriptomics to placental villous tissue from pregnancies complicated by PE and characterised important transcriptional changes in trophoblast and maternal cells^7–11^. In PE, EVTs have gene expression changes related to migration and cell death^8,9^, whereas STBs have changes related to angiogenic imbalance^7^ and senescence^11^. Among maternal cells impacted in PE are endothelial cells and immune cells, such as macrophages and T cells^7^. Moreover, Admati *et al*. quantified differences between Early PE (≤34 gestational weeks) and Late PE (>34 gestational weeks), confirming the heterogeneity of the disease^7^. In addition to placental dysfunction, recent evidence suggests aberrant transcription in other locations of the FMI such as the decidua basalis^11^, myometrial tissue^12^ and chorioamniotic membranes^13^. These studies often interrogate one tissue, missing the spatial heterogeneity of cell interactions within the FMI that underlie disease presentation and could reveal therapeutic targets.

Here, we focus on the severe presentation of the disease, which often involves maternal multi-organ dysfunction^14^ and preterm birth (<37 gestational weeks), with potential long-term health complications for both mother and fetus^3^. Despite the clinical and molecular importance of the placenta, elucidation of PE in its most florid state in the wider FMI is lacking. We present a spatially-resolved, multi-tissue single cell transcriptomic analysis of the entire FMI in a carefully phenotyped cohort of severe PE patients and gestational matched controls, spanning 25 to 37 gestational weeks. Correcting for gestational age, we quantify both suspected and novel fetal and maternal contributions to disease, per tissue and cell type, identifying distinct transcription underlying the Early extreme and Late moderate dysfunction in severe PE. In addition to concerted hypoxia, angiogenic imbalance, fibrosis and aberrant metabolism in the placenta, new maternal immune signatures, including changes in macrophages, mitochondrial dysfunction and interferon signalling across the FMI, offer an explanation to systemic inflammation and endothelial dysfunction in the mother, as well as new targets for therapy.

## Results

### Study cohort and multi-tissue sampling across the FMI

To study severe preeclampsia (PE) at single cell resolution across the entirety of the human FMI, we recruited a cohort of 20 donors, of whom 10 were experiencing severe PE from 25 to 37 weeks of gestational age, alongside 10 gestationally matched (± 1 week) controls. Hence, spanning across Early (≤34 weeks) and Late (>34 weeks) disease presentation (Fig. 1a and Extended Data Table 1). Gestation matched controls were from iatrogenic preterm deliveries without PE or any other maternal chronic disease, including diabetes. Donors were tested for presence of infection by standard clinical care but also with metagenomics (Supplementary Material and Extended Data Figure 1).

**Fig. 1:**
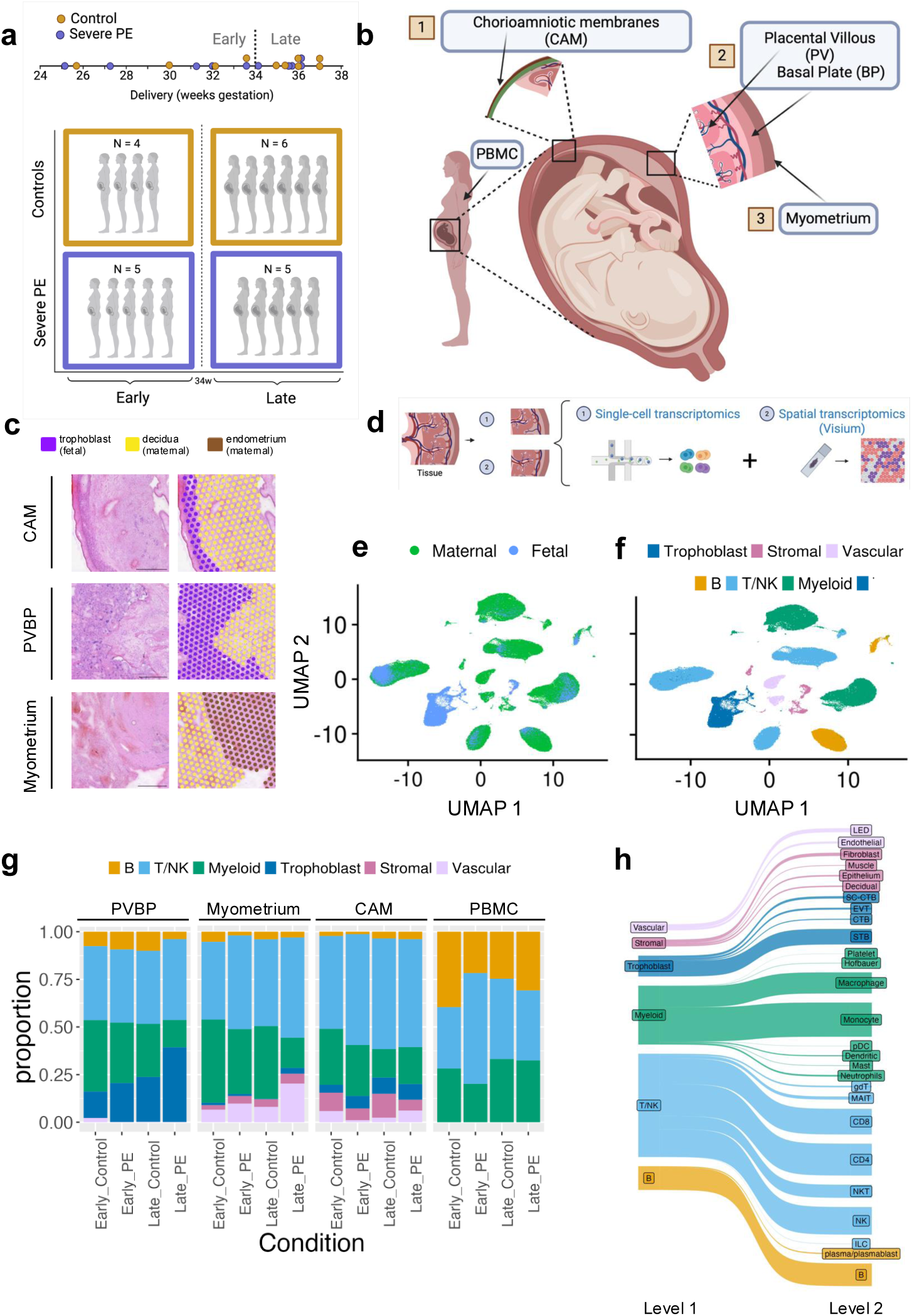
Multi-tissue single cell atlas of the human FMI with spatial context in severe PE. **a**, Cohort of 20 donors to study severe PE (N=10 cases and N=10 controls) with gestational ages from 25 weeks to 37 weeks across Early (≤34 weeks) and Late disease presentation (>34 weeks). **b**, Multi-tissue sampling per donor, including maternal blood (PBMCs), chorioamniotic membranes (CAMs), placental villi and adjacent basal plate (PVBP) and myometrium. **c**, Fetal and maternal populations sampling across the entirety of the FMI. Scale bars = 500 microns. **d**, Single cell transcriptomics and Visium spatial transcriptomics run on the same sample for each donor to spatially-resolve the effects of severe PE across the FMI. **e**, UMAP of fetal and maternal cells, identified using their genetic variation (SNPs) and trophoblasts markers. **f**, UMAP with the six major cell groups annotated (Level 1 annotation), including immune cell groups (B, T/NK and Myeloid) and non-immune cell groups (trophoblasts, stromal and vascular). **g,** These six groups were present across conditions. **h,** Clustering within each group from the Level 1 annotation resulted in annotations with increasing granularity, with 27 cell types in the Level 2 annotation. Level 3 annotation in Extended Data Figures 4,5.

We designed a systematic sampling strategy to profile cell populations from multiple tissues of the FMI for each donor. We sampled the placental villi and basal plate (PVBP), the chorioamniotic membranes (CAMs) and myometrium (Fig. 1b,c). This ensured the inclusion of fetal and maternal populations from the entirety of the FMI: placental villi (fetal), decidua basalis (maternal, from the basal plate and myometrium), chorion (fetal) and decidua parietalis (maternal, from the uterus). We also sampled peripheral maternal blood (PBMCs) at delivery per donor to study peripheral molecular signatures associated to disease.

### Constructing a multi-tissue single cell atlas of the FMI

To assess cell populations across the FMI, we performed single cell transcriptomics on the FMI tissues and PBMCs, without cell type enrichment (Fig. 1d). We analysed 292,773 high-quality cells (Extended Data Figure 2a). To classify cells as fetal or maternal, we leveraged their genetic variation^15^ as well as gene expression in fetal trophoblasts (Fig. 1e, Extended Data Figure 3). For donors and tissues, we manually annotated clusters based on differential gene expression and known markers. We first identified six major cell groups (Fig. 1f, Level 1 annotation), including immune cell groups (B, T/NK and Myeloid) and non-immune cell groups (trophoblasts, stromal and vascular) (Fig. 1g). We repeated the clustering in each group (Fig. 1h, Extended Data Figure 4), which resulted in 27 cell types (Level 2 annotation) and 59 cell types (Level 3 annotation) (Extended Data Figures 4,5). To complement this reference annotation, and provide spatial context to it, we performed Visium (10X Genomics) spatial transcriptomics on adjacent cuts from the same tissue sample (Figure 1b-d and Extended Data Figure 2). We used this FMI single cell atlas to inform the spatially resolved quantification of maternal and fetal cell contributions to severe PE.

### The importance of correcting for gestational age

We separately quantified the contribution of both gestational age and severe PE to molecular signatures of disease. We used the differential abundance of cell types as a proxy for this contribution, with unique abundance changes from gestational age (Late versus Early) and disease (severe PE versus control) revealing the impact of gestation and disease, respectively. The unique contribution of FMI tissues to severe PE is around 25% (Fig. 2a and Extended Data Figure 6), whereas it is only 10% in PBMCs. This agrees with severe PE being a placental disease, with the addition that cell contributions in the CAMs and myometrium are similar to those from the PVBP. A relevant but more modest contribution comes from peripheral tissues, as evidenced in PBMCs (Fig. 2a). Most molecular differences thus are impacted or confounded by gestation, even when cases and controls are matched within a week, underscoring taking gestational age into account in prenatal disease.

**Fig. 2:**
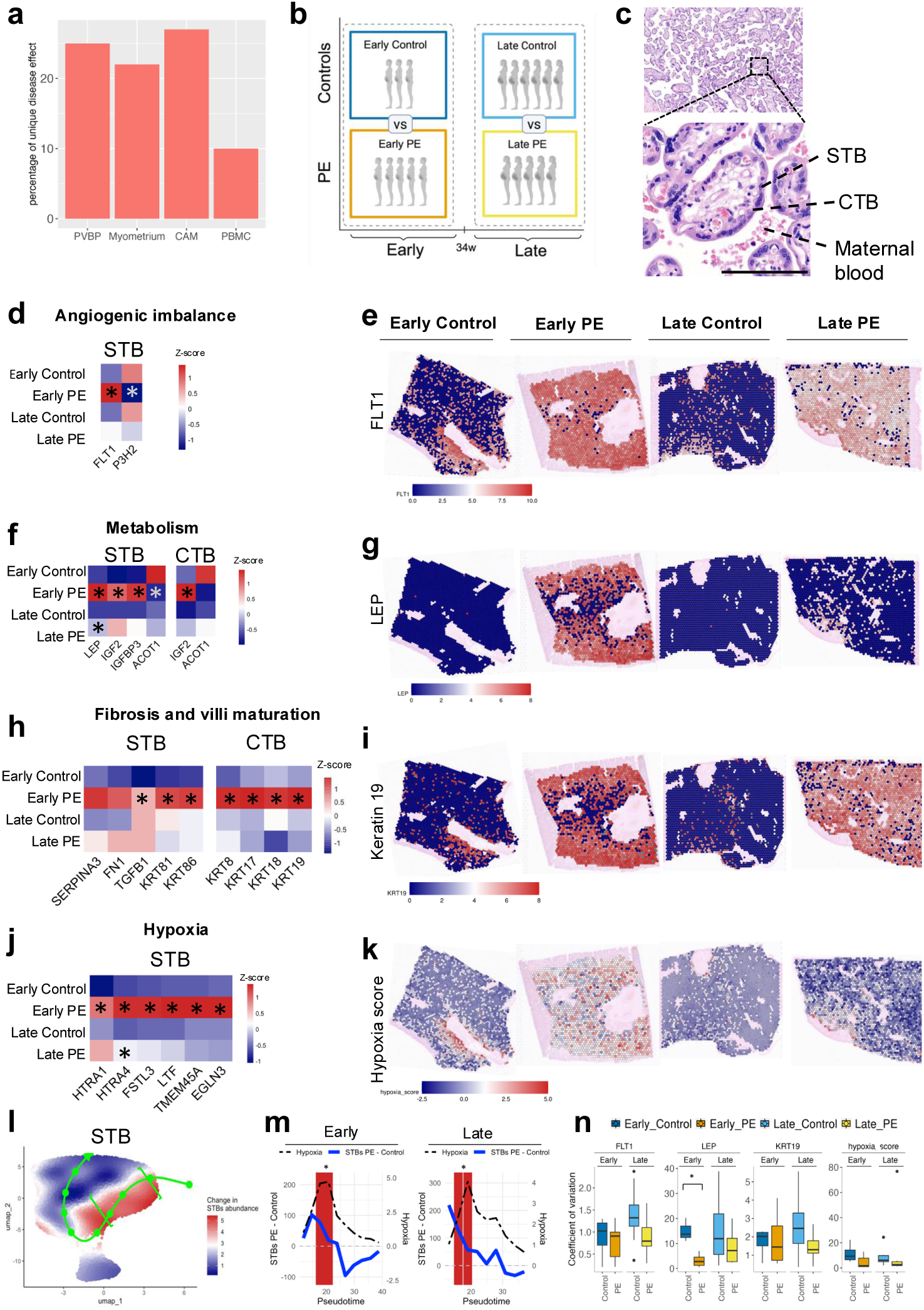
Severe PE involves angiogenesis imbalance, metabolic alterations and hypoxia of the placental villi. **a**, The unique contribution of severe PE to the molecular signatures per tissue (in %). **b**, Case-control experimental design considering gestational age and PE classification into Early and Late severe disease with a generalised linear model (GLM). **c**, H&E staining of the placenta villi with CTBs and STBs. Scale bar: 100 microns. **d,f,h,j**, Heatmaps of gene expression differences from our GLM. Expression is normalised and scaled in a Z-score across conditions per gene. * p<0.05. **d**, Genes involved in angiogenesis such as FLT1 and VEGFA are upregulated in Early PE. **e,g,i,k**, Gene expression across the tissue in representative samples per condition. Spatial plots with log-normalised counts. **e**, Spatial expression (Visium) of FTL1. **f**, LEP is upregulated in Early and Late PE in STBs while IGF2 is upregulated in both STBs and CTBs. **g**, Spatial expression of LEP. **h**, Genes associated to fibrosis such as TGFB1 and keratins are also upregulated in STBs and CTBs. **i**, Spatial expression of Keratin19. **j**, Heatmap of genes related to hypoxia on STBs. **k**, Spatial expression of the hypoxia score, combining the expression of hypoxia-related genes. **l**, UMAP of STB abundance between Early or Late PE and their matched controls along the STBs inferred pseudotime trajectory (green line). **m**, Difference in STB abundance (cell counts) in Early or Late disease versus matched controls, with significant differences (Wilcoxon rank-sum test) across pseudotime marked in light blue, where hypoxia scores are highest (dashed line). **n**, Variation in spatial expression across the placental villi for the significantly upregulated genes in e,g,i and k (*, Wilcoxon rank sum test, p<0.01).

We reached a comparable conclusion using gene expression as a proxy for the contribution of each cell type to gestation and disease. This is achieved with a generalised linear model (GLM), whose parameterisation quantifies Early and Late disease as well as the Average disease effect and Average gestational age (GA) effect (Extended Data Figure 6 and Supplementary Information). Importantly, this model identifies those cell types that contribute most to disease in severe PE, Early and Late (Fig. 2b and Extended Data Figure 6 d-g), which we discuss per tissue.

### Placental villi have spatially concerted pathological gene expression

We first examined the placental villi, fetal structures where gases and nutrients are exchanged between maternal and fetal blood (Fig. 2c). We found that STBs and CTBs (Fig. 2c, Extended Data Figure 5, Extended Data Figure 7a) contribute most to Early disease (Extended Data Figure 7c). STBs in Early PE have significantly higher expression of Fms related receptor tyrosine kinase 1 (*FLT1*), known to be clinically related to angiogenic imbalance^3^. Conversly Prolyl 3-Hydroxylase 2 (*P3H2*), whose activity is angiogenic, is significantly downregulated in Early PE^16^ (Fig. 2d, Extended Data Figure 7e). We also found overexpression of genes altering the metabolism. Leptin (*LEP*) was significantly upregulated in both Early and Late disease, consistent with previous reports^17,18^ (Fig. 2f, Extended Data Figure 7f). We also confirmed significant upregulation of Insulin-like growth factor-binding protein 3 (*IGFBP3*) in STBs and Insulin-like growth factor 2 (*IGF2*) in both STBs and CTBs^19^, alongside downregulation of Acyl-CoA thioesterase 1 (*ACOT1*), implicated in lipid metabolism^20^ (Fig. 2f).

Further, we found dysregulation of multiple keratins, mainly in Early disease. These include significant upregulation of *KRT18, KRT8* and *KRT19* in CTBs (Fig. 2h), which is associated with PE and implicated in trophoblast differentiation^21^. Interestingly, we find upregulation of additional keratins such as *KTR17* in CTBs and *KRT81* and *KRT86* in STBs (Fig. 2h). Keratin dysregulation has been associated with fibrosis in other tissues^22^. In addition, other genes promoting fibrosis and extracellular matrix deposition, such as *Transforming Growth Factor beta 1 (TGFB1), fibronectin-1* (*FN1*)^23^ and *SERPINA3*^24^ in STBs are overexpressed (Fig. 2h), suggesting that placental fibrosis broadly contributes to severe PE.

Finally, we identified upregulated genes in STBs, particularly in Early PE, increasing in hypoxia. These include *high temperature-requirement A4* (*HTRA4)* and *HTRA1*^25^, *Follistatin-like 3* (*FSTL3)*^26^, *Transmembrane protein 45A (TMEM45A)*^26^ and *Egl-9 Family hypoxia Inducible Factor 3 (EGLN3*)^27^ (Fig. 2j). We also found an STB subpopulation, overexpressing these hypoxic genes, that is unusually abundant in both Early and Late disease (Fig. 2l,m). They may underlie the ischemia and pathological increase of hypoxia in PE^28^.

We next investigated the spatial expression of these genes in the villi (Fig. 2, Extended Data Figure 7i,j), which validated their single cell expression (Fig. 2). In addition, we quantified the spatial variation of genes overexpressed in severe PE and found homogeneous gene expression across the villous region. This supports a concerted and shared stress response of trophoblasts from the villous tree (Fig. 2n), which is accentuated in Early disease but also present in its Late presentation.

### Extravillous trophoblasts in severe PE change in abundance at the decidua basalis

Although defective spiral artery remodelling by the fetal extravillous trophoblast (EVT) has been implicated in PE^2,3,6^, it is unclear whether this is due to decreased density or depth in the invasion of EVTs, with immunohistochemistry providing contradictory interpretations^29,30^. We evaluated this by defining spatial domains as anatomical regions in a tissue, sharing histology and gene expression. In the PVBP, we distinguished two domains: the placental villi, of fetal origin, and the decidua basalis, of maternal origin and proximal to the villi (Fig. 3a). Similarly, we separated the Myometrium into the decidua basalis (distal to the villi) and muscle (Fig. 3c), both maternal.

**Fig. 3:**
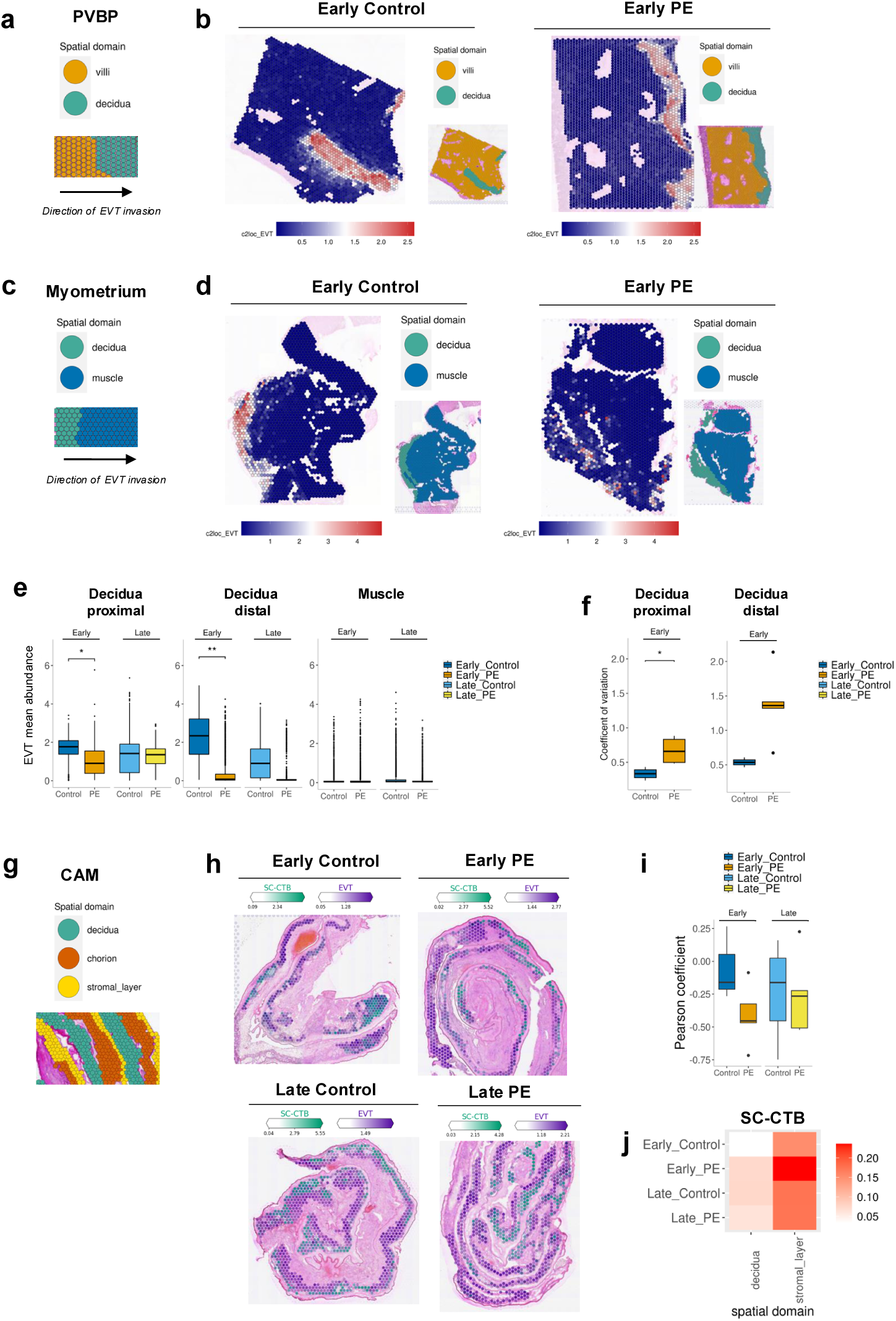
Extraplacental trophoblast populations have a distinct spatial distribution at the FMI. **a**, Spatial domains at the PVBP are the placental villi and the adjacent decidua basalis. **b**, Deconvoluted abundance of extravillous trophoblasts (EVTs) at the decidua basalis (PVBP). **c**, Spatial domains at the Myometrium are the decidua basalis and muscle. **d**, Deconvoluted abundance of EVTs at the Myometrium. **e**, The spatial abundance of EVTs in the decidua basalis is reduced in Early PE compared to Early controls, particularly at the distal end (*p < 0.05, ** p < 0.01, mixed-effects model). **f**, Coefficient of variation of spatial EVT abundance across the decidua basalis, proximal and distal to the villi in the PVBP, with significant variation in Early (p < 0.01) (Wilcoxon rank sum exact test **g**, Spatial domains at the CAM are the maternal decidua parietalis, the fetal chorion and the fetal stromal layer (within a rolled CAM). **h**, SC-CTBs and EVTs coloured by their abundance in the chorion. **i**, Pearson correlation coefficient of SC-CTBs and EVTs location in the chorion, which is negative suggesting that they are spatially anticorrelated. **j**, Neighbouring spots of those with high SC-CTB abundance, expressed as a proportion. The SC-CTBs are closer to the stromal layer than the decidua.

We detected a significant decrease of EVTs in Early PE in the decidua basalis, proximal to the villi (p < 0.05) (Fig. 3b,e) but particularly distal to the villi (p < 0.01) (Fig. 3d,e), pointing to a decreased depth of invasion. We also quantified the spatial distribution of these EVTs and found that they are sparsely distributed across the decidua basalis (Fig. 3f). This is though less evident in Late disease. Finally, some EVTs are present in the muscle in the Myometrium but their numbers do not allow for statistical interpretation (Fig. 3e). Thus, failure of spiral artery remodelling by EVTs in severe PE is from both reduced density and depth of invasion^29^.

### Extraplacental trophoblasts in the decidua parietalis are spatially patterned

The FMI also extends to the CAM, where we defined spatial domains of fetal and maternal origin (Fig. 3g). In the chorion, we identified EVTs that are transcriptionally similar to those at the decidua basalis, as previously reported^31^. We also confirmed a recently described population of trophoblast, the SC-CTBs, characterised by expression of keratins *KRT6A, KRT14, KRT17* and integrin subunit *ITGB6*^31^ (Extended Data Figure 5). Interestingly, we noticed that the tissue distribution of EVTs and SC-CTBs is spatially anticorrelated (Fig. 3h,i), particularly in Early PE. This is the result of SC-CTBs in Early disease favouring the stromal layer (Fig. 3j), stratifying their distribution in the CAM.

### Numerous maternal immune cell types have mitochondrial dysfunction

Inflammation and alterations in several maternal immune cell types have been reported in severe PE^32^. We found, in Early disease, upregulation of genes involved in mitochondria function and oxidative stress across maternal immune cell types in the PVBP, Myometrium and CAM (Fig. 4a, Extended Data Figure 8a). These are genes involved in the production of reactive oxygen species (ROS) (several subunits of complex I nicotinamide adenine dinucleotide and hydrogen dehydrogenase: MT-ND1, MT-ND2, MT-ND3, MT-ND4, MT-ND4L, MT-ND6), *cytochrome c oxidase subunit I and 2* (MT-CO1, MT-CO2), *cytochrome b* (MT-CYB) and *mitochondrial ATP synthase complex* (MT-ATP6, MT-ATP8). Affected cell types include both innate cell types (NK, NKT and macrophages) and adaptative immune cell types (CD8 and CD4 T Cells). Dendric cell types (pDC, cDC1, cDC2 and cDC3) are at low numbers in our analysis, yet they also present mitochondria alterations. This indicates broad immune maternal mitochondrial dysfunction across the FMI in Early PE. Interestingly, this may also be the case in some fetal immune populations (Extended Data Figure 8c).

**Fig. 4:**
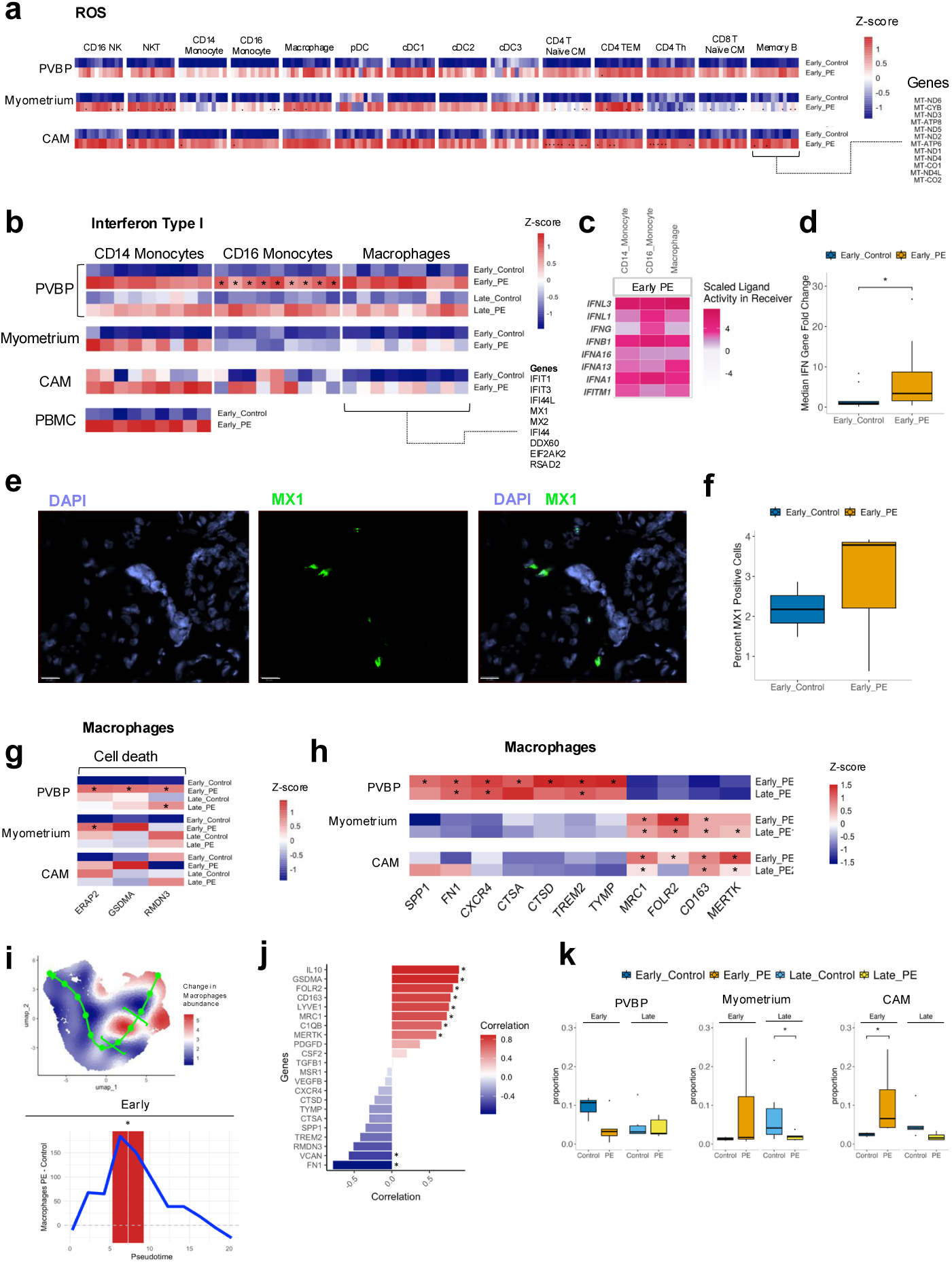
Maternal immune effects in severe PE across the FMI. **a**, Mitochondrial genes involved in oxidative stress genes are upregulated in multiple cell types in Early disease. Gene expression is normalised and scaled using Z-score across conditions per gene (*p<0.05, GLM). **b**, Interferon type I stimulated genes (ISG) are upregulated in several myeloid cell types. **c**, Ligand-receptor activity supports the activation of IFN-I. **d**, qPCR of ISGs in PBMCs, with significant upregulation in Early disease (*p <0.05, Wilcoxon Rank Sum). **e**, Example of immunohistochemistry in the PVBP in Early PE of an ISG (MX1, green) and DAPI (blue). Scale bars=20 microns. **f**, Immunofluorescence quantification of MX1 positive cells in the PVBP for two cases and three controls in Early disease. **g**, Macrophages upregulate genes related to cell death. **h**, Macrophages in severe PE have gene expression differences across the FMI. Gene expression is normalised and scaled using Z-score across tissues per gene. (*p < 0.05). **i**, Top panel: UMAP of changes in macrophage abundance in PE against controls along the inferred macrophage pseudotime trajectory (green line). The pseudotime interval considered for downstream analysis is in brackets. bottom panel: difference of cell counts in Early disease (blue line). The red background indicates pseudotime intervals where the cell counts difference is significant (Wilcoxon rank-sum test, *p < 0.05. **j**, Correlation between the gene expression and cell abundance along pseudotime colored by significance from a Person’s correlation test. **k**, Proportion of macrophage in Early and Late disease per tissue (*p<0.05).

### Type I interferon is upregulated in the myeloid compartment

We also identified enrichment of interferon stimulated genes (ISGs) in maternal CD14 Monocytes, CD16 Monocytes (p <0.05) and Macrophages in the PVBP in both Early and Late disease (Fig. 4b, Extended Data Figure 8b). Ligand-receptor activity of ISGs also supports an enhanced IFN type 1 response in the myeloid compartment (Fig. 4c). Moreover, MX1 immunofluorescence (representative ISG) supports upregulation of IFN-1 (Fig. 4e,f). Notably, there is upregulation of ISG in CD14 Monocytes also in the Myometrium, CAM and PBMCs, although it did not reach significance (Fig. 4b). We thus corroborated it in maternal circulation in Early disease by qPCR (Fig. 4d). Thus, there is a pathological type I interferon response in the myeloid compartment across the FMI, one that is detectable in maternal circulation within 25 – 34 weeks of gestation. Of note is the complementary, significant upregulation of IFITM3 in fetal monocytes as well (Extended Data Figure 8d).

### Macrophages at the PVBP upregulate genes associated with cell death and fibrosis

We then investigated maternal macrophage responses at the FMI. This highly heterogenous cell type has been implicated in pregnancy complications, including PE^7,33^. We identified, with our GLM, significant upregulation of *Gasdermin A* (*GSDMA*) in Early disease at the PVBP (Fig. 4g), with also elevated expression in the Myometrium and CAMs (Fig. 4g). *GSDMA* is a pore-forming protein in the mitochondrial membrane involved in pyroptotic cell death^34^. Two other genes whose overexpression leads to apoptosis, the mitochondrial *RMDN3 (PTPIP51)*^35^ and *ERAP2*^36^, are also significantly upregulated in Early PE at the PVBP (Fig. 4g). In agreement, macrophages in this tissue are reduced (Fig. 4k), possibly from selective cell death.

Further, the anti-inflammatory molecule *TREM2* expression is upregulated in Early^7^ and Late PE in the PVBP (Fig. 4h). Macrophages in the PVBP also overexpress several proteases and scavenger proteins, including Cathepsins (*CTSD, CTSA*) and *MSR1 (*CD204)^37^, as well as pro-fibrotic *SPP1/osteopontin*, *FN1* and *CXCR4*^38–40^ (Fig. 4h). This suggests that macrophages in the PVBP are impacted and likely contribute to placental fibrosis, a cardinal feature of Early PE.

### Macrophages at the myometrium and CAM upregulate phagocytic and anti-inflammatory genes

Meanwhile, macrophages at the myometrium and CAM have similar gene expression (Fig. 4h), including significant upregulation of phagocytosis genes *MRC1* (CD206), and anti-inflammation genes *CD163, FOLR2* and *MERTK*^38,41,42^ in both Early and Late PE. There is also a subpopulation of macrophages in the Myometrium and CAM (Fig. 4i,j) upregulating additional anti-inflammatory genes, such as *IL10*, and *LYVE1* (Fig. 4j)^41,42^. Towards this end, macrophages expand in Early PE in the Myometrium and CAM, the opposite behaviour of macrophages in the PVBP (Fig. 4k). Altogether, these suggest that macrophages are distinctly activated across the FMI in disease. They decrease through apoptosis in the PVBP, while promoting fibrosis, and expand in the Myometrium and CAM to fight inflammation. These are hallmarks of lost homeostasis in these tissues under severe PE.

### Long-range effects of trophoblasts in severe PE

Some of the products of the upregulated genes we identified in STBs, for example *LEP, IGF2* and *FLT1* (Fig. 2), have been identified in maternal circulation^3,18^, perhaps contributing to the systemic pathophysiology of severe PE. We thus assessed cell-to-cell communication from STBs to other cell types, jointly in the PVBP and Myometrium. We identified the *LEP* receptor, *LEPR*, highly expressed in monocytes and macrophages as well as in endothelial and NK maternal cells in both Early and Late PE. Notably, *LEPR* is also expressed in STBs suggesting a positive feedback loop enhancing *LEP* effects (Fig. 5a). We also identified TGFB1 receptors (produced by STBs) in some myeloid and endothelial cells (Extended Figure 8i). In the endothelial cells of the myometrium, we also detected downregulation of genes associated with angiogenesis (Extended Data Figure 8h). So, in addition to the well-known systemic endothelial dysfunction effect of FLT1^3^, other long-rage effects from the villi to myeloid and endothelial cell types may contribute to the severe PE presentation.

**Fig. 5:**
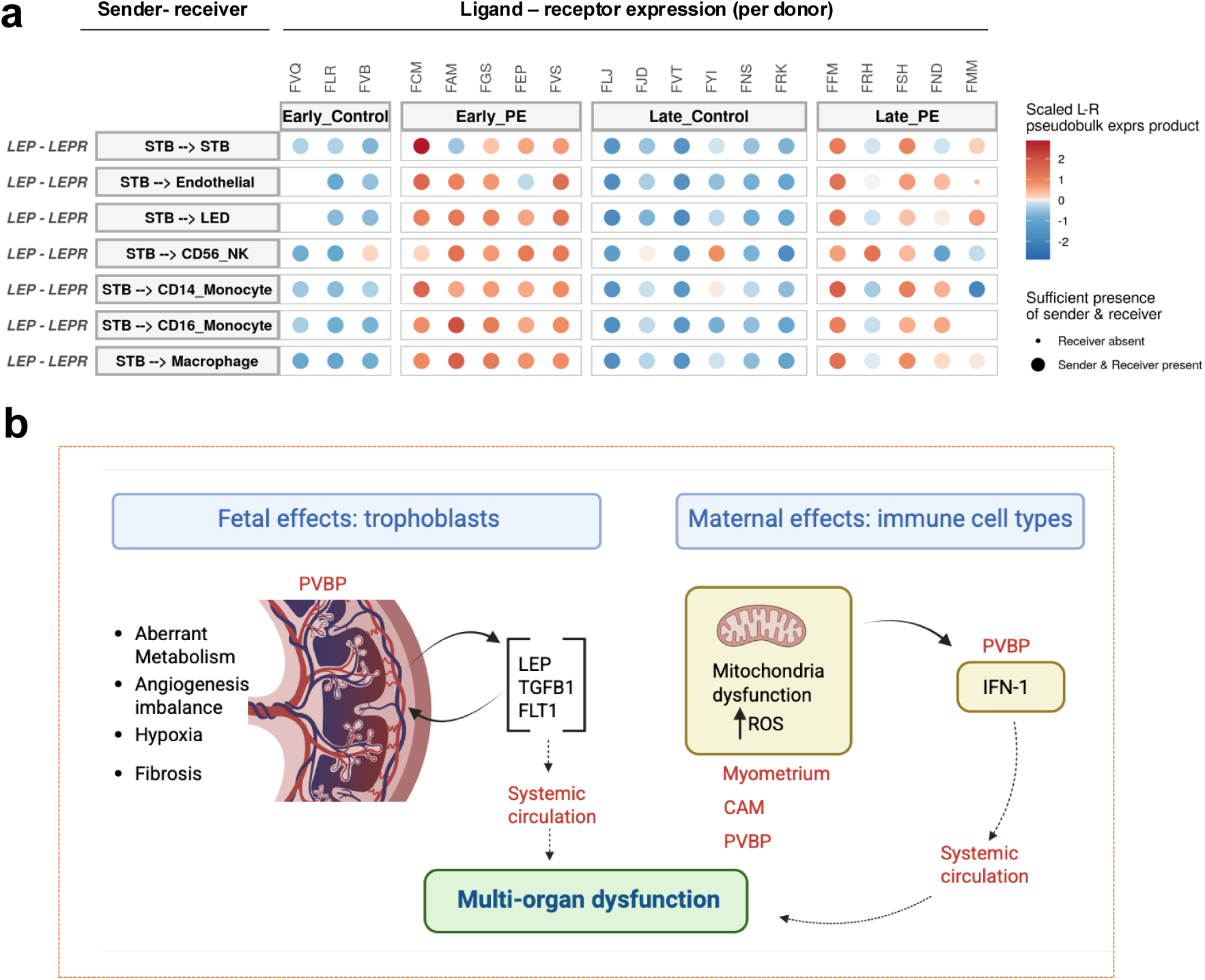
Trophoblasts long-range effects might impact maternal cell types in severe PE. **a**, Ligand-receptor communication, per donor, between STBs expressing LEP and immune cell types expressing its receptor LEPR. Interactions with a prioritization score above 0.5 are shown. Dot plot depicts scaled ligand-receptor pseudobulk expression product. **b**, Fetal effects from the villous tree include the release of molecules to systemic circulation, impacting multiple organs. Among the maternal effects in immune cell types, we uncover mitochondria dysfunction and interferon type I expression. We detect IFN-I in maternal circulation, further contributing to the systemic and multi-organ pathology of severe PE.

## Discussion

Sampling all the tissues from the FMI, while correcting for gestational age, allowed us to identify the distinct fetal and maternal molecular contributions to severe PE. We spatially and quantitatively characterised them, revealing broad dysregulation of the placental villi, where a combination of angiogenic, metabolic, fibrotic and hypoxic fetal alterations accompany the disease (Fig. 5b). We confirmed the reduced invasiveness of EVTs in severe PE^2^, which becomes more pronounced far from the villous region, amplifying the impact of fetal dysfunction. In addition, LEP is strongly upregulated at the placental villi in Early and Late disease. This is important as leptin induced ROS, unfolded protein response and ER stress, as well vascular endothelial dysfunction in female mice^17,18^. In this context, our inferred LEP cell-to-cell communication within STBs may enhance placental dysfunction. In addition, LEP communication to endothelial and immune cell types in the adjacent decidua and myometrium, together with the reported leptin increases in maternal serum of PE^18^, may cause systemic effects (Fig. 5b). Leptin antagonists have shown promise in autoimmune diseases^43^ and are an exciting avenue for STB targeted therapies.

We reveal among the maternal contributions to disease, a novel interferon type I response in the myeloid compartment, along with mitochondria oxidative stress in numerous immune cell types in the myometrium, CAM and PVBP in Early disease (Fig. 5b). This is in addition to the known mitochondria dysfunction and oxidative stress in STBs^44^. This maternal oxidative stress is implicated in the activation of IFN-I in several immune conditions with systemic inflammation, such as systemic lupus erythematous (SLE)^45^ and juvenile dermatomyositis^46^. Notably, patients with SLE are at an increase risk of developing PE and have upregulated IFN signalling that contributes to vascular damage at the FMI^47^. Elevated IFN-I also resulted in abnormal EVT migration in an *in vitro* system^48^. We thus propose IFN-I as an unrecognised contributor to systemic endothelial dysfunction in severe PE. Its presence in peripheral blood in severe PE, also demonstrated here, promises better screening of patients that might benefit from anti-interferon^49^ or antioxidant therapeutics^46,50^.

Further to ectopic IFN-I in the diseased myeloid compartment, we also unravelled maternal responses in macrophages across tissues at the FMI with severe PE. At the PVBP, macrophages express pro-fibrotic molecules that may contribute to the placental fibrosis observed in histology. The hypoxic tissue environment and IFN/mitochondrial stress can trigger selective apoptosis^51^, decreasing the response to local inflammation despite the expression of anti-inflammatory molecules such as TREM2^7^. In contrast, myometrial and CAM macrophages express a molecular signature, including MERTK, MRC, LYVE, that in rheumatoid arthritis (RA) potentially resolves inflammation and contributes to remission^42^. As new cell type-specific treatments emerge, we advance that some macrophage subpopulations may be a therapeutic target.

In conclusion, we present novel mechanistic insights into severe PE, a disease orphan of treatment, opening different tissue and cell-specific responses for therapeutic intervention. Given the severity of molecular dysfunctions in Early disease, compared to its Late presentation, their timely consideration during gestation is likely beneficial and could change the extremely poor prognosis of severe PE.

## Methods

### Donor inclusion and exclusion criteria

We recruited pregnant women at the fetal medicine unit, antenatal care unit and labour ward of the University College London Hospitals, NHS Foundation Trust. The study cohort comprised 20 pregnant women, 10 of which diagnosed with severe preeclampsia (PE) and 10 gestational matching controls ranging from 25 to 37 weeks gestational age across Early (<34 weeks) and Late (>34 weeks) disease presentation. Severe PE is defined as the presence of blood pressure more than or equal to 160/110 mmHg, accompanied by significant proteinuria (more than or equal to 300 mg protein/day) with haematological or biochemical changes or severe maternal symptoms. The study was approved by the South Oxford Research Ethics Committee (REC CODE: 17/SC/0432).

Inclusion criteria were women with a singleton pregnancy, able to consent, experiencing preterm delivery by C-section due to iatrogenic reasons or severe PE prior to 37 weeks of gestation. Exclusion criteria were women with pregnancies affected by major fetal anomalies (whether chromosomal, structural, or genetic), twin pregnancies and comorbidities including diabetes mellitus (gestational or T1 or T2), maternal autoimmune, renal or heart disease. Donors were excluded if had any clinical symptoms/signs of infection or a positive test of infection during standard clinical practice. In addition, we ruled out possible undetected infections using metagenomics. Our metagenomics analysis identified an *Ureaplasma* pathogenic bacteria in donor FJJ, which was then excluded from our comparative analysis.

### Tissue sampling

Placenta Villi and Basal Plate (PVBP) samples were collected from the mid-point of the largest distance between the cord insertion site and the edge of the placenta. Myometrial samples were taken via biopsy from the area of the placental bed mirroring the expected placental sampling-site. From the edge of the placenta closest to the placental tissue sampling site chorioamniotic membranes (CAMs) were cut from the placental edge and rolled, with the end closest to the placenta being in the middle of the roll. For each tissue, adjacent samples were collected, one was placed in cold HypoThermosol FRS for dissociation and single cell transcriptomics, and another was snap frozen and embedded in OCT medium for spatial transcriptomics. Up to 24 hours prior to delivery 7ml of peripheral venous maternal blood was collected for PBMC isolation. Further details can be found in the Supplementary Information.

#### Metagenomics

PVBP and CAM samples for each donor were processed including an additional patient (FCK) which had clinical signs of infection and acted as a positive control. Parallel DNA and RNA extractions using the ZymoBIOMICS™ DNA/RNA Miniprep Kit were performed. RNA and DNA libraries were prepared as previously described for Illumina sequencing^52^ with microbial enrichment using NEBNext® Microbiome DNA Enrichment Kit following the manufacturers protocol but using double the amount of MBD2-Fc-bound magnetic beads per nanogram (ng). Enriched DNA was processed with the NEBNext® Ultra™ II FS DNA Library Prep Kit for Illumina. RNA libraries were prepared using the KAPA RNA HyperPrep Kit with RiboErase. RNA and DNA libraries from each batch were sequenced either on the Illumina Nextseq 2000 (Pilot and Batch 1) or Illumina Novaseq 6000 (Batches 2 and 3) to achieve between 20M-50M reads per sample. These were demultiplexed and ran through the CZID web-based platform for taxonomic profiling^53^ and filtering using the package Metathresholds^52^.

### Single cell tissue dissociation

PVBP, CAM and Myometrial samples were dissociated on a gentleMACS Octo dissociator in 10ml Accutase (PVBP and CAM) or 5ml Accumax (Myometrium) using a custom protocol (see Supplementary Material). Dissociated cell suspensions were then passed through stacked 100um and 70um cell strainers and subjected to red cell lysis using Red Cell Lysis buffer (Milteny Biotec). PVBP and CAM samples were then subjected to dead cell removal using Dead Cell Removal MicroBeads loaded onto MACS LS columns (Miltenyi Biotec). Myometrial samples were pelleted and resuspended in 1x DPBS, 0.04% BSA and passed through a 40um Flowmi Cell Strainer (Sigma-Aldrich). In addition, nuclei single cell suspensions were prepared for the CAM from donors “FVQ”, “FCM”, “FGC”, “FLJ”, “FRK”. Briefly, snap frozen and OCT embedded CAM samples were cryosectioned and nuclei were isolated from the sections using Chromium Nuclei Isolation Kit (10X Genomics), following 10X Genomics user guide CG000505. PBMCs from peripheral venous maternal blood were isolated using Histopaque 1077 (Sigma-Aldrich).

### Single cell and spatial transcriptomic library preparation and sequencing

Dissociated cells, isolated PBMCs and isolated nuclei were checked for concentration and viability using an Acridine Orange/Propidium Iodide Stain (logos Biosystems) on a logos Biosystems Luna FL automated cell counter. Single cell suspensions per tissue and donor were used to generate single cell/nuclei transcriptomic and single cell/nuclei TCR libraries. We loaded 20,000 cells/nuclei into a 10X Genomics Chromium Controller using the Chromium Next GEM Chip K and Chromium Next GEM Single Cell 5’ Kit v2 kits (10X Genomics) as per 10X Genomics user guide CG000331.

Ten-micrometre sections of snap frozen PVBP, Myometrium and CAM samples were loaded onto Visium Spatial Gene Expression slides as recommended by 10X Genomics. The area of the PVBP samples chosen for loading onto the relevant capture areas contained the edge proximal to the decidua basalis. Loaded sections on Visium slides were fixed, stained and imaged on a Motic EasyScan One slide scanner. Spatial transcriptomic libraries were prepared following manufacturer guidelines. Permeabilization times of 18 minutes for CAM and Myometrium and 6 minutes for PVBP, were established as appropriate using the Visium Spatial Tissue Optimisation kit (10X Genomics).

Resulting spatial transcriptomic, single cell and single nuclei transcriptomic and TCR libraries were sequenced on an Illumina Novaseq 6000 using five S4 (200 cycle) v1.5 sequencing kits (Illumina), with a configuration of Read 1: 28 cycles, Index read 1: 10 cycles, Index read 2: 10 cycles, Read 2: 190 cycles. Libraries were sequenced to a minimum coverage of 20,000 reads per cell or 25,000 reads per Visium spot.

### Immunofluorescence assay

PVBP samples from Early PE Early Controls were cryosectioned and loaded onto one glass slide. Sections were fixed in 10% Formaldehyde and then blocked and permeabilized using 1X DPBS 0.1% Tween 10%FBS 0.1% Triton-X with 10mg/ml dextran sulphate for 60 minutes. Overnight staining was performed using 1X DPBS 0.1% Tween 10% FBS with 10mg/ml dextran sulphate and a 1:100 dilution of CoraLite® Plus 488-conjugated MX1 Recombinant antibody (Proteintech). Then nuclear stain DAPI was added at 5ug/ml for 1 minute and a coverslip was then added using SlowFade Gold Antifade mounting medium. Stained slides were imaged on a Nikon Ti2 inverted microscope at 20X magnification. Resulting image files were loaded into QuPath 0.5.0. Morphological regions were delineated to include an approximately equivalent proportions of decidua and villous placenta to capture both maternal and fetal contributions. Positive cell detection analysis was performed using DAPI for nuclei detection and a nuclear, maximum intensity threshold for detection of MX1 positive cells.

### Quantification of IFN-I expression in PBMCs

An extended cohort of Early PE (N= 14) and gestational matched controls (N=9) was used to measure IFN-I expression in PBMC (see details on Supplementary Material, Extended data table 2). PBMCs isolated from whole blood were subjected to RNA extraction using the Qiagen RNEasy plus kit. Extracts where then subjected to qPCR using taqman probes for 4 interferon related genes (IFIT1, IFI44L, ISG15, RSAD2) along with 1 housekeeping gene (18S). The NEB Luna® Universal Probe One-Step RT-qPCR Kit was used and qPCR reactions were performed using an Applied Biosystems Quantstudio 5 Real-Time PCR instrument, using cycling conditions as per recommended for the reaction kit. Output files were processed using the Design & Analysis Software by Applied Biosystems with Ct values per replicate group exported as a .csv file. DDCt values were calculated and then converted into fold change values as per 2^DDCT. The median value of the above four interferon related genes per sample was calculated. The Wilcoxon test was used to compare cases and controls for each gene, along with the median interferon gene fold change.

### Alignment, count matrix and ambient correction of scRNA-seq/snRNA-seq data

For each single cell and single nucleus library, the raw bcl files were converted into fastq files using mkfastq (version cellranger-7.1.0; 10X Genomics). Sequencing reads were aligned to the GRCh38-2020-A human reference genome (GENCODE v32/Ensembl98 distributed by 10X Genomics) and a matrix of unique molecular identifier (UMI) counts and TCR clonotypes per library was obtained using cellranger “multi”. We used CellBender 0.2.0 to correct for ambient RNA per library and classify cell-containing droplets from empty ones^54^.

### Fetal and maternal cell classification in single-cell transcriptomics

For cell classification into fetal or maternal, we used *Freemuxlet* v.1 (popscle software)^15^. First, a variant site list (VCF) with the most common SNPs from the 1000 Genomes Project was used as a panel of SNP positions. Then, the *dsc-pileup* function was used to identify the allelic composition, base quality, and the number of reads from each allele at each common SNP location per library using the BAM file output from cellranger. We combined the dsc-pileup outputs from the tissue PVBP with Myometrium or PBMCs or CAM per patient to circumvent the low abundance or absence of fetal cells in the Myometrium and PBMC and run Freemuxlet using two groups (*i.e* two individuals: *--nsample 2*) and default parameters.

To identify the inferred genotype (0, 1 or 0/1) that corresponded to the fetus or mother or doublet, we calculated the average expression of a set of trophoblast markers (“CGA”, “CYP19A1”, “GH2”, “PAPPA”, “VGLL1”, “PAPPA2”, “HLA-G”) per genotype. We assigned as fetal the genotype with higher average trophoblasts expression. To verify the Freemuxlet genotype assignment, we used a DNA microarray (Infinium Global Diversity Array v1.0) on a subset of samples (8 pairs of fetus and maternal donors using DNA isolated from cord blood and PBMC respectively).

### Single cell Quality Control (QC)

Doublets were detected in two ways: 1) using Scrublet 0.2.3^55^ (*manual threshold = 0.25*) per library; and 2) as a droplet with a mixed genotype (0/1) from Freemuxlet. These were filtered out. Seurat 4.2.1^56^ was used to calculate QC metrics. Cells were filtered out if: 1) number of detected genes was below 400 and above 6,000; 2) number of UMIs was above 30,000; 3) percentage of mitochondrial reads was more than 12% and 4) percentage of haemoglobin genes was more than 0.25.

### Single cell integration and cell type annotation

After pre-processing, gene expression matrices from all donors and all tissues were integrated using Seurat 4.2.1^56^. Briefly, a standard workflow including normalisation (“LogNormalise”), scale (“ScaleData”) and highly variable genes selection (2,000 genes) were used to perform principal component analysis (*npcs = 25*), k-nearest-neighbour calculation (*k.param = 20*) and graph-based community detection using Louvain clustering. Harmony 0.1.0^57^ was used to correct for batch effects from the single cell gene expression and single nuclei gene expression methods differences.

We annotated cell types using gene expression of manually curated genes from the literature and genes differentially expressed per cluster (obtained using Seurat function “FindMarkers”). Cell types were annotated at three levels of resolution (refereed as annotation level 1, level 2 and level 3). First, a coarse cell group annotation rendered six major groups: 1) Trophoblasts; 2) Stromal; 3) Vascular; 4) Myeloid; 5) B; and 6) T/NK cells at annotation level 1. Subsetting each of these groups and repeating the clustering pipeline described above allowed the assignment of two more granular levels of annotation. During clustering, we detected further doublets (identified by a mixed profile of gene expression in the cluster) and those were also filtered out.

### Cell abundance differences from disease and gestational age

To measure the differential abundance of cell populations at single-cell resolution due to disease or gestational age, we used dawnn 1.0.8^58^. Using as an input a Seurat object per tissue, we calculated a KNN graph using the first 10 principal components (*reduced_dim = pca*) for the PVBP, Myometrium and PBMCs tissues. For the tissue CAM, we used Harmony 0.1.0^57^ (*reduced_dim = harmony)* to correct for batch effects introduced by the two different single cell methods and used the first 10 components from harmony. We applied a false discovery rate correction with the Benjamini-Yekutieli procedure as implemented in dawnn 1.0.8 using an *alpha = 0.05*.

We tested for differential abundance separately comparing: 1) conditions (Control or PE); or 2) gestational age (Early or Late). We measured whether a cell is called differentially abundant in one comparison, in both or in neither. Donor FJJ was excluded of this and the rest of the comparative analysis as explained above.

### Cell type differential gene expression analysis per tissue

We tested for single cell differential gene expression in each tissue while accounting for gestational effects and donor-to-donor variability. Cell types with less than 25 cells per condition or present in only one donor were excluded from the differential expression analysis. Genes lowly expressed (less than 20 total counts) were also filtered out.

For each tissue, we first aggregated the measured counts per donor and cell type at annotation level 3 into a pseudobulk expression profile. We use EdgeR to fit a log-linear model using the design matrix: ∼0 + *c*, where *c* corresponds to the condition:

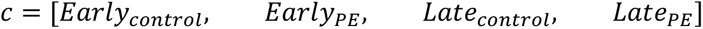

We defined Early disease as the contrast: *Early_PE_ vs Early_control_* and Late disease as: *Late_PE_ vs Late_control_*

We tested for differential expression per contrast using a likelihood ratio test (LRT) implemented in EdgeR 3.36.0^59^. Multiple testing was controlled by using a Benjamini-Hochberg (BH) correction. Genes were considered differentially expressed if the logFC >0.5 and FDR <0.05.

### Macrophages differential expression across tissues

To test for differences in gene expression of Macrophages across tissues we also used a GLM framework (EdgeR). Selecting only PE Macrophages (Early PE or Late PE), we modelled gene expression using a paired design to adjust for donor-specific effects using the design matrix: ∼0 + *tissue* + *donor*. As before, we used a log-linear model for tissues *k* = 3 and the contrasts were set to compare Macrophages from *PVBP vs Myometrium* and *PVBP vs CAM*.

### Cell proportions per tissue

To test for differences in cell proportions accounting for donor variability and gestational age, we used the GLM framework implemented in the *propeller* function of the R package speckle 0.0.3^60^. We use a matrix with donors *d* as rows and conditions *c* as columns to fit a linear model using the design matrix: ∼0 + *c*.

For *tissue* = {*CAM*}, we added a term to correct for the effect introduced by using two different single-cell methods using the modified design matrix: ∼0 + *c* + *method*. We tested for differences in cell proportions per cell type using a moderate t statistical test per contrast and applied a false discovery rate correction using BH procedure.

### Differential abundance and gene expression in pseudotime in STB and Macrophages

We used *slingshot v.2.6.0*^61^ to infer cell trajectories in pseudotime following the Condiments pipeline^62^ to identify pseudotime intervals (as a proxy of cell subpopulations) containing differential abundance of cells of any condition (control or severe PE). We used in-house R scripts to generate a sequence of pseudotime intervals of *n* pseudotime units (by dividing the maximum pseudotime by 10 to obtain *n* intervals). For each interval, we obtained the number of cells from each donor or the proportion of cells relative to the donor’s total number of cells. To test for difference in cell counts (or proportions) in Early or Late disease, we used a Wilcoxon test followed by a Benjamini-Hochberg correction for multiple testing (threshold 0.2) performed in intervals with cells from at least 3 donors from each condition.

To correlate shifts in gene expression with shifts in cell abundance across pseudotime intervals, we calculated the mean gene expression of a given gene (or set of genes) per cell at each pseudotime interval and correlated this expression with the difference between the mean of cells per donor in severe PE cases compared to controls in each interval.

### Cell-to-cell communication

*MultiNicheNet v1.0.3*^63^ was used for differential ligand-receptor analysis in Early and Late disease. Cell types from the tissues PVBP and Myometrium were integrated into a single Seurat object to take advantage of their spatial continuity. Cell types at annotation level 3 with less than 25 cells and coming from one single donor were excluded. Nichenet v2 ligand-receptor network and matrices from “*organism = human*” were obtained from Zenodo (DOI: 10.5281/zenodo.7074291). Early disease and Late disease were modelled using the same GLM as before. We applied a p-value cutoff of 0.05 and log-fold change threshold of 0.5. Other parameters were set as default. We selected the top 150 targets per ligand and visualise the top predictions using MultiNicheNet inbuilt functions.

### Alignment and gene count per spot of Visium

Demultiplexing, alignment of reads to the human reference genome (GRCh38-2020-A, GENCODE v32/Ensembl98 distributed by 10X Genomics), identification of spots for each tissue and quantification of gene counts per spot were performed using spaceranger v2.0.0 (10X Genomics).

### Spatial domain identification per tissue

Spatial domains were identified manually by two researchers using the 10X Genomics Loupe Browser 7. Annotations were cross referenced to generate a consensus. First, a *kmeans* clustering of the gene expression per capture area (*k=4* for CAM and *k=3* for PVBP and Myometrium tissues) was used to generate clusters which broadly aligned with the morphologically expected tissue domains. A combination of differential expression analysis (in-built in Loupe browser) and knowledge of cell type markers with known spatial distribution (i.e. decidual cells in the decidua basalis and decidua parietalis from PVBP and CAM, respectively) was used to manually annotate spatial domains. Poor quality spots, artifacts or low-confidence spatial domain assignment were manually labelled as “background” and were excluded from further analysis.

### Spatial hypoxia gene expression score in placenta villi

The hypoxia score included the genes: “HTRA1”, “HTRA4”, “FSTL3”, “EGLN3”, “TMEM454” and was calculated using the Seurat function *AddModuleScore*^56^ using 30 control genes on spots belonging to the spatial domain “villi” per capture area.

### Cell type deconvolution

*cell2location v0.1.3*^64^ was used to deconvolute cell type abundances per spot in Visium. We trained a single cell regression model for 350 epochs using the cell type annotation at level 2 as a single cell reference (excluding donor FJJ), keeping genes expressed in at least 3% of the cells and excluding genes expressed in less than 5 cells. To deconvolute the cell types per spot, we use *N_cells_per_location = 8* and a *detection_alpha = 20.* We trained the deconvolution model for 3,000 epochs and exported the 5% quantile of posterior distribution for visualisation. Each Visium capture area was deconvoluted separately.

### Spatial expression statistics using a mixed-effects model

To test for differences in abundance of EVT per spot in Early or Late disease, we used a linear mixed-effects model implemented in the packages *lme4 1.1-27.1* and *lmerTest 3.1-3*. We considered the condition (Early Control vs Early PE or Late Control vs Late PE) as the fixed effect while the variation between donors belonging to the same condition was considered a random effect using the model:

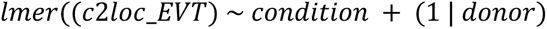

### Spatial correlation of EVT and SC-CTB in CAM

A Pearson correlation was used to quantify spatial correlation of EVT and SC-CTB in the CAM spatial domain “chorion”. We filtered out spots with SC-CTB abundance less than 0.5. For statistical comparison between conditions in Early or Late disease, we used a Fisher’s Z-transformation and computed a p-value (two-tailed test).

### Neighbourhood analysis in the CAM

To quantify the spatial arrangement of spots with high abundance of the cell type SC-CTB (abundance greater than 3) in the spatial domain “chorion” (CAM), we calculated the identity of their neighbour spots using the nearest neighbour algorithm (*k=2*) as implemented in the *RANN 2.6.1* package.

## Acknowledgements

We thank Sasha Tunnock, Tyrrell Hatch and Claire Boevink, research midwives at University College London Hospitals, NHS Foundation Trust. Also, Dale Moulding from the Light Microscopy Core Facility at UCL Great Ormond Street Institute of Child Health, and Ciaran Hutchinson and Olumide Ogunbiyi from the GOSH Histopathology Department. Finally we would like to thank UCL Genomics for providing the sequencing of single cell and spatial transcriptomic libraries.

ECR is funded by a senior research fellowship awarded by the Kennedy Trust for Rheumatology Research (KENN 21 22 09) and a research prize from the Lister Institute for Preventive Medicine. This work was supported by the MRC (MR/W028158) and the NIHR GOSH Biomedical Research Center (NIHR203326).

## Author Contributions

YESC developed analytical strategies, performed data analysis and contributed to data interpretation and writing of the manuscript. TX designed the and performed the experiments and contributed to data interpretation and writing of the manuscript. YESC and TX contributed equally to this work. JJMV performed genotyping and pseudotime analysis and contributed to writing of the manuscript. LF performed metagenomic experiments and data analysis and contributed to writing of the manuscript. NJS, ECR and LRW provided advice and reviewed the manuscript. SC conceived of and supervised the project, contributed to data interpretation and writing of the manuscript. SLH conceived of and supervised the project and contributed to data interpretation and writing of the manuscript. SC and SLH jointly supervised the work.

## Data availability

Single cell and Visium libraries will be deposited in a public repository.

## Code availability

The code repository to reproduce analysis steps included in this manuscript can be found at https://github.com/sanchezy/FMI_severePE/.

---

**Extended Data Figure 1.**
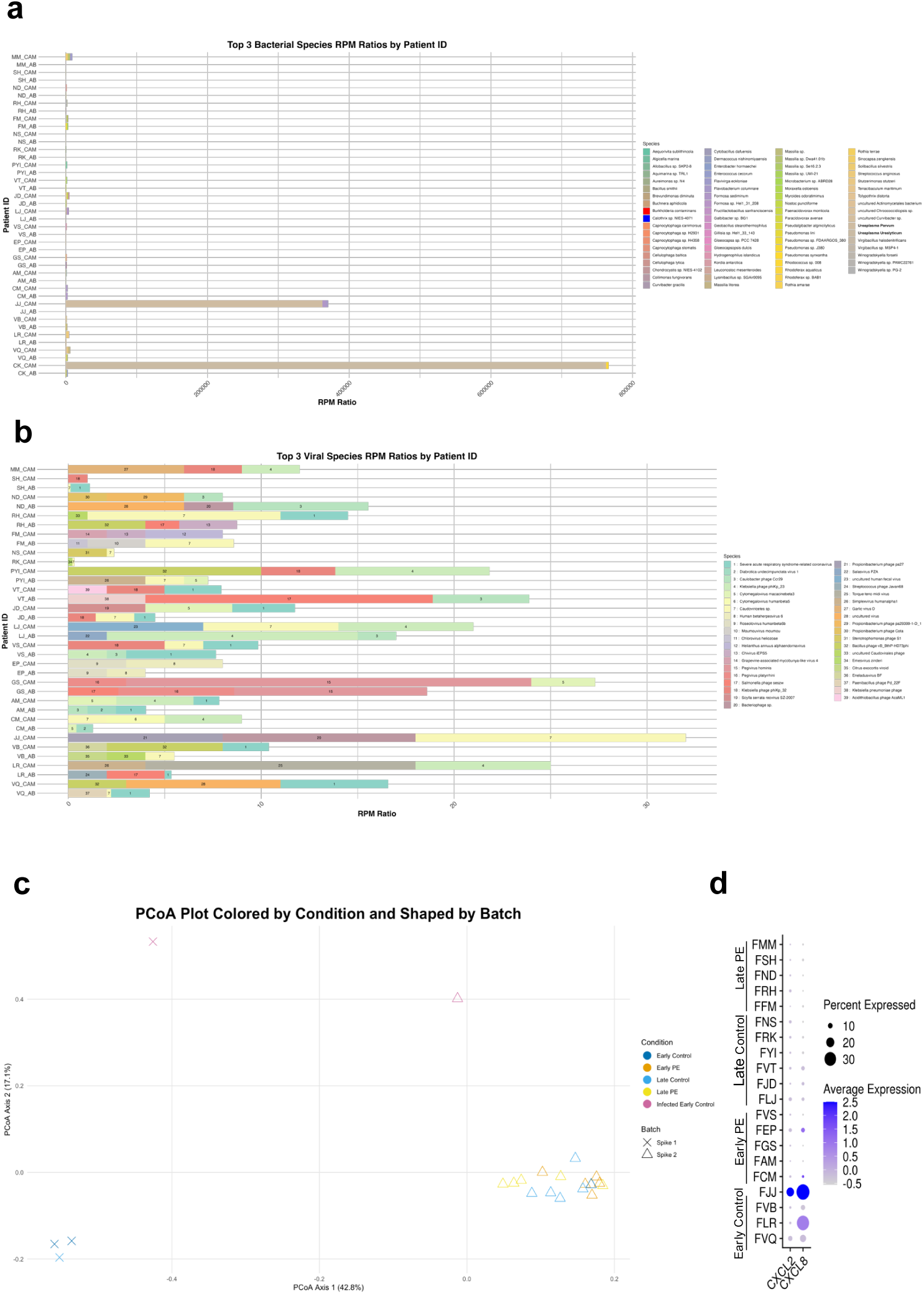
Metagenomic analysis of CAM and PVBP to screen for infection. **a,** Ratio of reads per million (RPM) for each species relative to the negative control of the top three bacterial species (x-axis) for each sample (y-axis). Samples include PVBP (AB) and chorioamniotic membranes (CAM) tissue per patient with combined DNA and RNA results for each tissue type. Where the same species is present in both RNA and DNA samples for one sample type, the highest RPM ratio was used. A known-infected (non-PE) sample (CK) was run alongside every batch as a form of external positive control. **b,** RPM ratios of the top three viral species (x-axis) for each sample (y-axis). **c,** Bray-curtis Principal Coordinate Analysis (PCoA) of microbial proportions for each patient where all CAM and PVBP data are combined per patient. PE condition indicated by colour. **d,** CXCL2 and CXCL8 expression differing for FJJ sample in the spatial domain decidua parietalis of the CAM (from Visium).

**Extended Data Figure 2.**
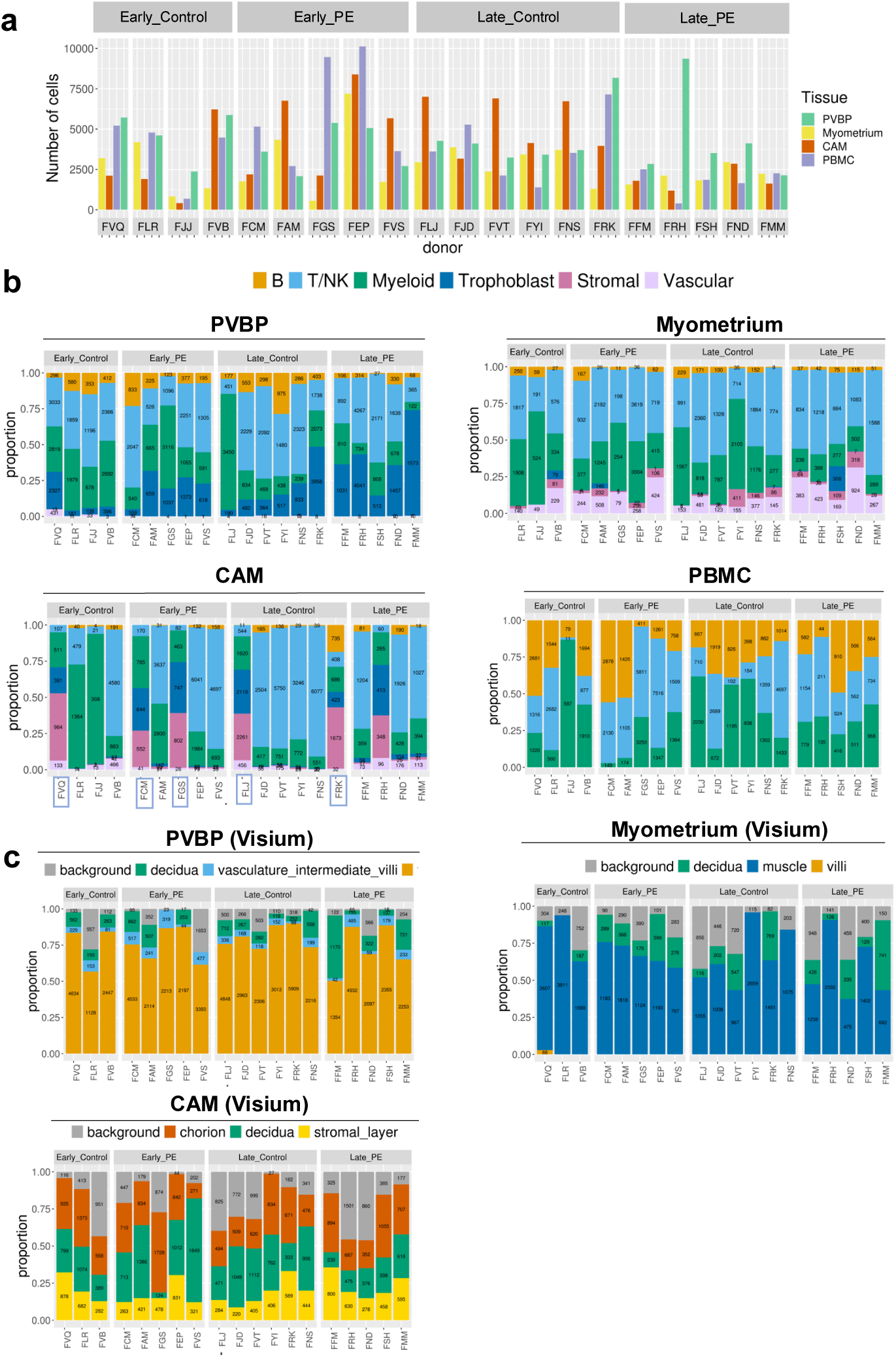
Single cell transcriptomics and spatial transcriptomics sampling per donor. **a,** Number of cells that have passed quality control per tissue and per donor. **b,** Proportion of sampled cell groups (annotation level 1) per tissue and per donor. For CAM tissue, donors that were profiled using single-nuclei gene expression are indicated with blue rectangles. **c,** Proportion of spots per spatial domain sampled for each tissue and donor using Visium 10X Genomics. PVBP included 25 Visium capture areas, Myometrium included 20 capture areas and CAM included 19 capture areas. PVBP, placenta villous basal plate. CAM, chorioamniotic membranes. PBMC, peripheral blood mononuclear cells.

**Extended Data Figure 3.**
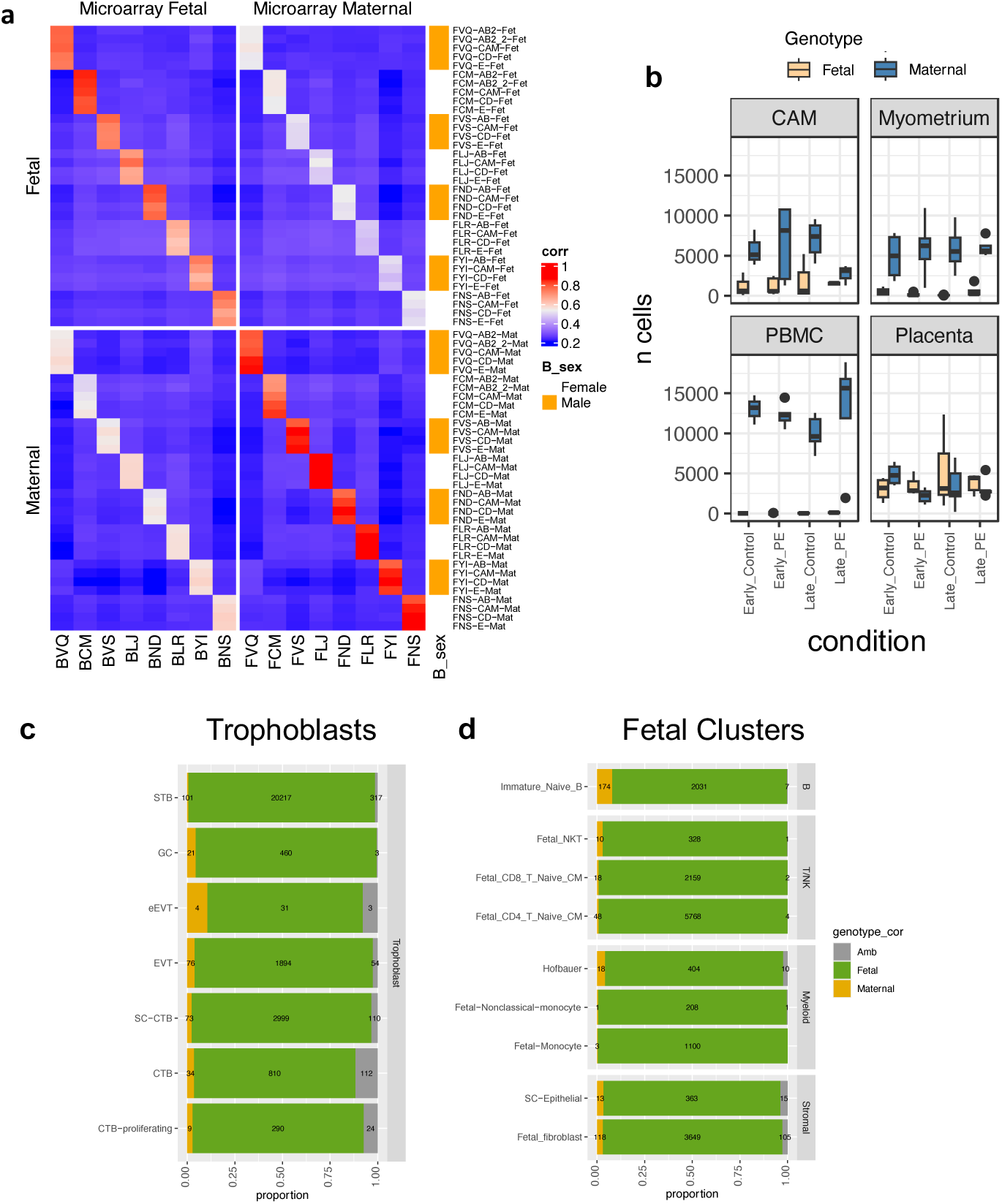
Cells demultiplexed into fetal or maternal origin from their genetic variation. **a,** Heatmap with Pearson correlation coefficients for each Freemuxlet-cluster per library (x axis) and microarray SNP genotypes (y axis) based on allelic composition. Libraries from pregnancies with male babies are annotated in orange. Labels in x and y axes indicate patient pair (fetal-maternal pair). In the x axis, labels are followed by tissue type (AB: PVBP; CAM; Chorio-amniotic membrane; CD: Myometrium; E; maternal PBMCs). **b,** Number of cells classified as maternal or fetal origin by tissue type and condition. **c,** Proportion of cells classified as fetal, maternal or ambiguous Trophoblasts. These cells are from fetal origin and its classification as maternal and ambiguous is considered erroreous, which is under 3.5% of cells. **d,** Proportion of fetal, maternal or ambiguous cells classified by freemuxlet in other fetal cell clusters (defined fetal from the graph-based clusters containing more than 80% cells classified as fetal by freemuxlet).

**Extended Data Figure 4.**
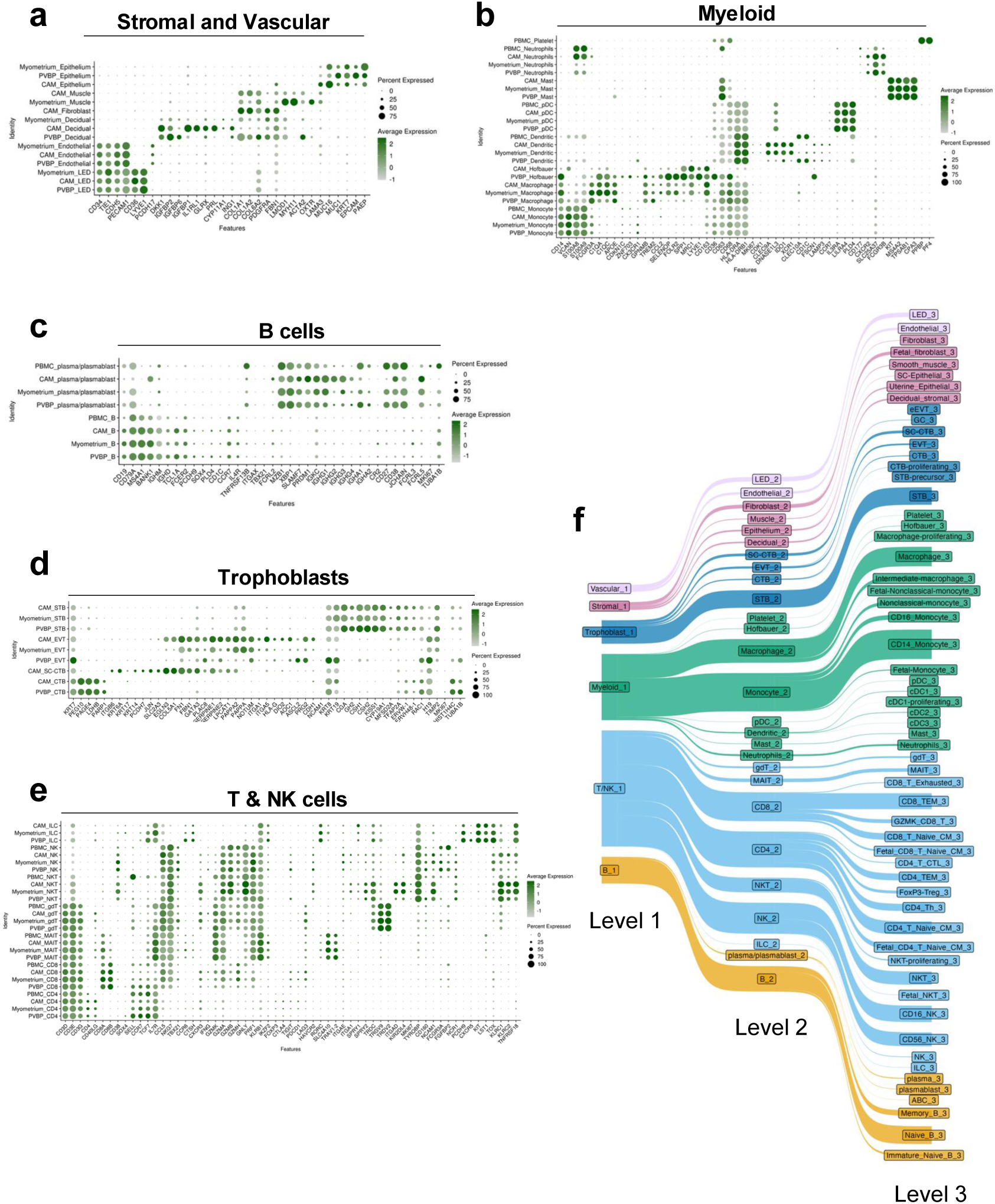
Marker gene expression used for Level 2 manual annotation. **a-e,** Dot plots of markers used to define five cell groups at level 2 of annotation per tissue. **a,** Stromal and Vascular. **b**, Myeloid. **c,** B cells. **d,** Trophoblasts. **e,** T and NK cells. **f,** Sankey plot depicting three levels of annotations. PVBP, placenta villous basal plate. CAM, chorioamniotic membranes. PBMC, peripheral blood mononuclear cells.

**Extended Data Figure 5.**
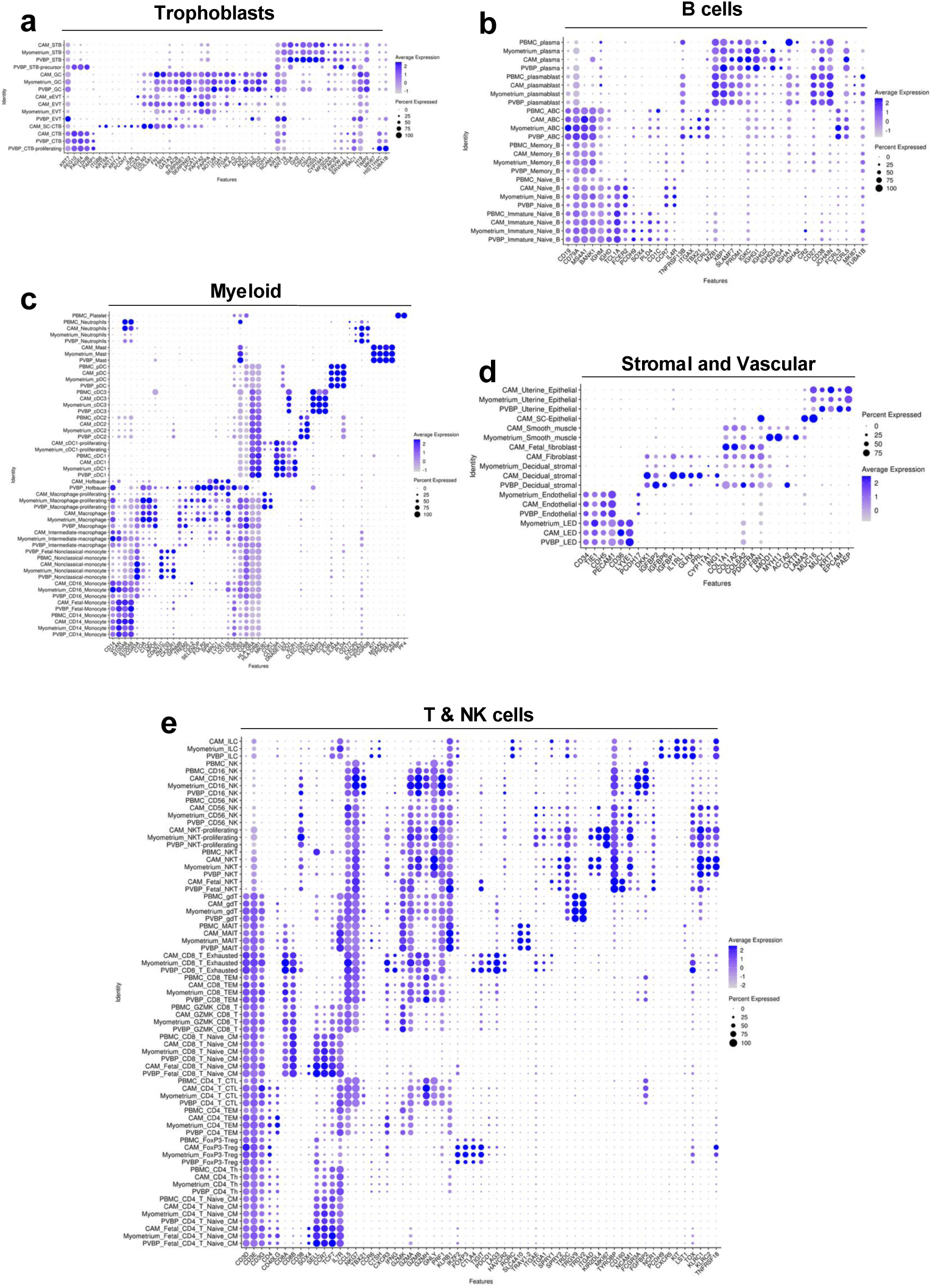
Marker gene expression used for Level 3 manual annotation. **a-e.** Dot plots of markers used to define five cell groups at level 3 of annotation per tissue. **a,** Trophoblasts. **b**, B cells. **c,** Myeloid. **d,** Stromal and Vascular. **e,** T and NK cells.

**Extended Data Figure 6.**
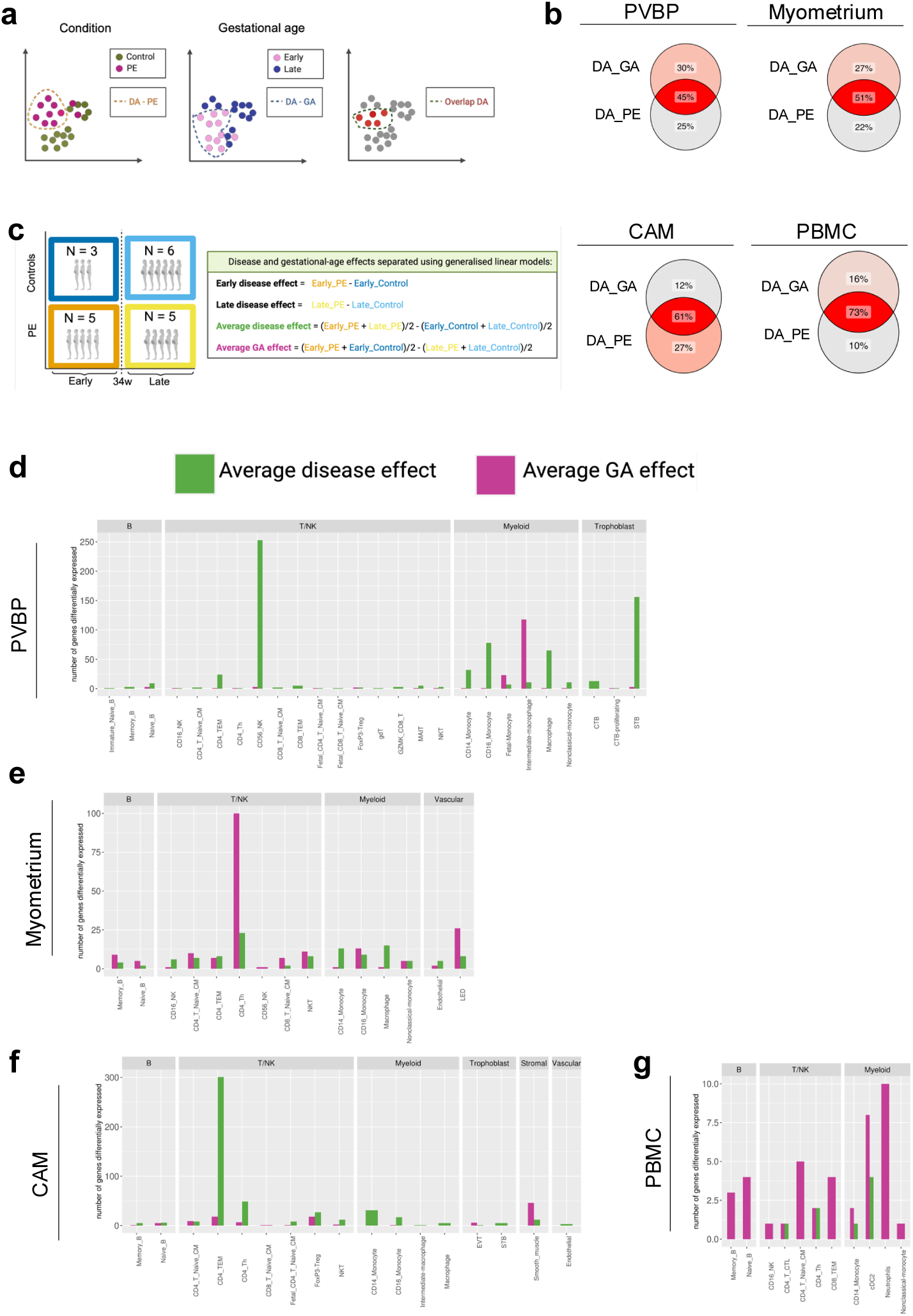
Gestational age and disease effects can be separated using generalised-linear models. **a,** Schematics of the single-cell differential abundance (DA) analysis taking as a variable the condition (Control and PE) or gestational age (Early and Late), separately. The overlap DA are the cells that are called differential abundant on both comparisons. **b,** Venn diagrams showing the percentage of differential abundant cells due to gestational age (DA_GA), due to disease (DA_PE) or the overlap. **c,** The gestational age effect and disease effects can be separated using generalised linear models (GLM) considering four groups: Early Control, Early PE, Late Control and Late PE. This parameterisation allows for quantification of Early and Late disease as well as the Average disease effect and Average gestational age (GA) effect. **d-g,** Number of differentially expressed genes (DEG) per cell type as a proxy of the disease or gestational age in different tissues. Notably, cell types affected in disease are not affected due to gestational age and vice versa. Only cell types with DEG are shown. DEG were filtered by p <0.05 and log Fold Change > 0.05.

**Extended Data Figure 7.**
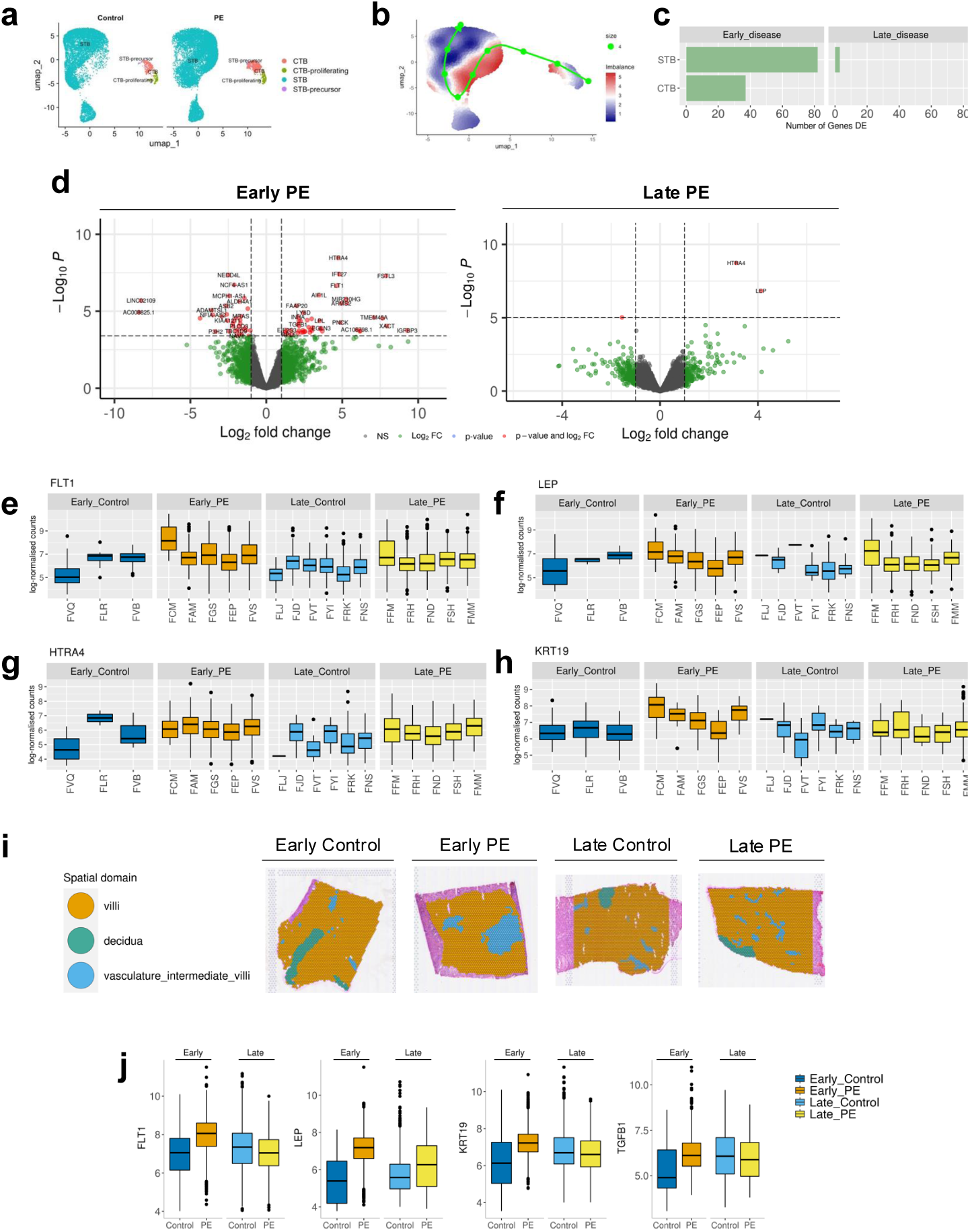
Severe PE single cell and spatial characterisation of trophoblasts in the placental villi. **a,** UMAP showing trophoblast populations, coloured by cell type and separated by condition at the villi. **b,** UMAP displaying the abundance difference between Early Control or Late Control and Early PE and Late PE. Pseudotime trajectory (green line) is initialised in the cell type CTB-proliferating. **c,** Number of differentially expressed genes in STBs and CTBs (p adjusted value below 0.05 and log fold change above 0.5). **d**, Volcano plots of STBs in Early and Late disease. **e-h,** Selected overexpressed genes plotted per donor in STBs (log normalised counts). **e,** FLT1, **f**, LEP, **g**, HTRA4, **h**, KRT19. **i,** Spatial domains in the PVBP are villi, decidua and vasculature intermediate villi. Representative Visium images of Early Control, Early PE, Late Control and Late PE. **j,** Gene expression in the spatial domain villi of selected genes (log normalised counts).

**Extended Data Figure 8.**
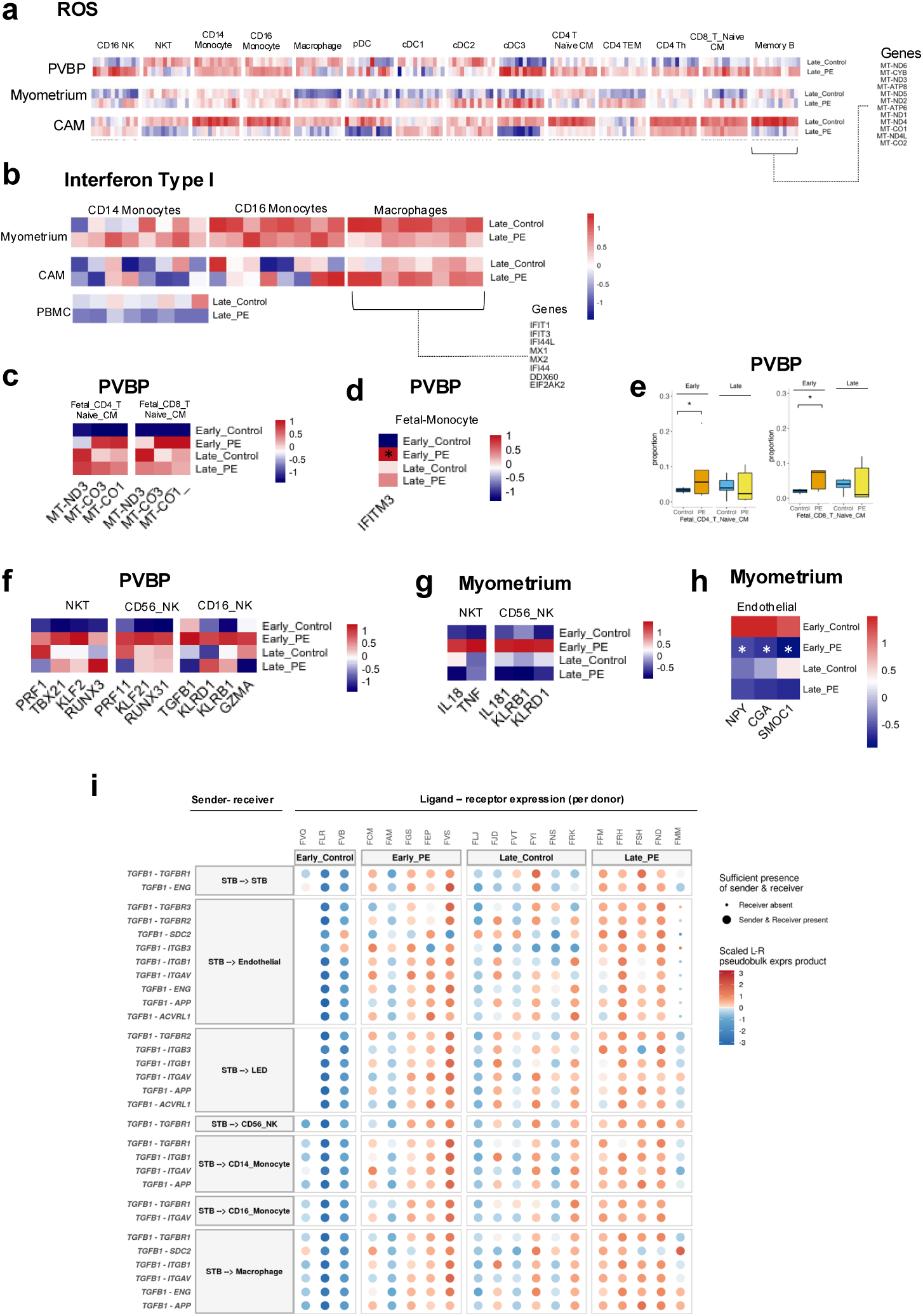
Maternal cell signatures in severe PE. **a,** Heatmap of mitochondrial genes in several cell types in Late disease. Expression is normalised and scaled in a Z-score across conditions per gene. **b,** Heatmap of interferon type I expression in some myeloid cell types in Late disease. **c**, Heatmap of mitochondrial genes in the Fetal T CD4 Naive/CM and Fetal T CD8 Naive/CM in the tissue PVBP. **d,** Heatmap of IFITM3 gene in fetal monocytes. **e**, Boxplots of cell proportions of Fetal T cells per condition. **f,** Some activation markers in NKT, CD56 NK and CD16 NK cells in the PVBP tend to be upregulated in disease. **g**, Some activation markers of NKT and CD56 NK in the Myometrium. **h,** Endothelial genes involved in angiogenesis are downregulated in the Myometrium (p<0.06, GLM). **i**, Ligand-receptor inferred communication between trophoblast expressing TGFB1 (STB) and immune cell types expressing its receptor per donor. Interactions with a prioritization score above above 0.5 are shown. Dot plot depicts scaled ligand-receptor pseudobulk expression product.

## Extended Data tables

Table 1. Severe PE cohort description (Excel file)

Table 2. IFN-I cohort description (Excel file)

Table 3. Differential expression in Early PE (csv file)

Table 4. Differential expression in Late PE (csv file)

