## Supplementary material for "Spatially resolved fetal and maternal cell contributions to severe Preeclampsia"

1. Prospective cohort design
   1.1. Patient inclusion and exclusion criteria
   1.2. Clinical definition of severe preeclampsia (PE)
2. Tissue sampling
   2.1. Placenta villi and basal plate (PVBP)
   2.2. Myometrium
   2.3. Choriamniotic membranes (CAMs)
   2.4. Maternal Peripheral Blood Mononuclear Cells (PBMCs)
3. Metagenomics
   3.1. Nucleic Acids extraction
   3.2. Enrichment and library preparation
   3.3. Sequencing
   3.4. Analysis
4. Single cell and spatial transcriptomics experiments
   4.1. Single cell tissue dissociation
   4.2. Single cell library preparation
   4.3. Spatial transcriptomics library preparation
   4.4. Sequencing
5. Immunofluorescence assay
   5.1. Sectioning and fixation
   5.2. Blocking, permeabilization and staining
   5.3. Imaging and positive cell quantification
6. Quantification of IFN-I expression in PBMCs
7. Single cell transcriptomics analysis
   7.1. Pre-processing
   7.2. Fetal and maternal single cell classification
   7.3. Single cell Quality Control (QC)
   7.4. Single cell integration and cell type annotation
   7.5. Cell abundance differences from disease and gestational age
   7.6. Cell type differential expression analysis per tissue
   7.7. Macrophages differential expression across tissues
   7.8. Cell proportions per tissue
   7.9. Differential abundance and gene expression in pseudotime
   7.10. Cell-to-cell communication
8. Spatial transcriptomics analysis
   8.1. Alignment and gene counts per spot

8.2. Spatial domain identification per tissue
8.3. Spatial hypoxia gene expression score in placenta villi
8.4. Cell type deconvolution
8.5. Spatial expression statistics using a mixed-effects model
8.6. Spatial correlation of EVT and SC-CTB in CAM
8.7. Neighbourhood analysis in the CAM

#### 1. Prospective cohort design

##### 1.1 Patient inclusion and exclusion criteria

We recruited pregnant women at the fetal medicine unit, antenatal care unit and labour ward of the University College London Hospitals, NHS Foundation Trust. The study cohort comprised 20 pregnant women, 10 of which diagnosed with severe preeclampsia (PE). The study was approved by the South Oxford Research Ethics Committee (REC CODE: 17/SC/0432). All participants signed informed consent, voluntarily gifting samples to the study and were free to withdraw from it.

Our inclusion criteria were women with a singleton pregnancy, able to consent, experiencing a threatened preterm labour or diagnosed with severe PE prior to 37 weeks of gestation. Our exclusion criteria were women with pregnancies affected by major fetal anomalies (whether chromosomal, structural, or genetic), twin pregnancies and comorbidities including diabetes mellitus (gestational or T1 or T2), maternal autoimmune, renal or heart disease. In addition, for each participant signs of infection (viral or bacterial) were investigated, and patients were excluded if any clinical symptoms/signs were identified (including increased white cell count or c-reactive protein, increased temperature in conjunction with either high HR and /or low blood pressure). Participants were also excluded if we received infection positive urine, vaginal or blood samples, taken as part of clinical care. In addition, we ruled possible undetected infections using metagenomics (as described in section **3. Metagenomics**). Our metagenomics analysis identified an *Ureaplasma* pathogenic bacteria in donor FJJ, which was then excluded from our cohort.

The following maternal characteristics were recorded for each donor: age, ethnicity, body mass index, parity (prior to delivery), gestation of disease onset, and administration of antenatal corticosteroids within one week of delivery. In the severe preeclamptic cohort, further information taken from medical records included highest blood pressure prior to delivery, treatment initiated, serum biochemical abnormalities and urine protein-creatinine ratio. The following fetal/neonatal characteristics were recorded: abnormal fetal dopplers, the presence of intrauterine growth restriction (<1$^{st}$ centile), estimated fetal weight centile (using Hadlock algorithm), birth weight (gm) and sex of the neonate (see **Extended Data Table 1** for details).

##### 1.2 Clinical definition of severe preeclampsia (PE)

Severe PE is defined as the presence of blood pressure more than or equal to 160/110 mmHg, accompanied by significant proteinuria (more than or equal to 300 mg protein/day) with haematological or biochemical changes or severe maternal symptoms [1]. PE is stratified into early (before and including 34 weeks of gestational age) and late onset (more than 34 weeks of gestational age) [2].

#### 2. Tissue sampling

All sampling occurred within 30 minutes of delivery to ensure RNA integrity. Below, we describe sampling per tissue type.

##### 2.1 Placenta villi and basal plate (PVBP).

The placental tissue was washed twice in 1x DPBS (-Ca, -Mg). From the mid-point of the largest distance between the cord insertion site and the edge of the placenta, one sample was taken from each donor and two adjacent 2 x 2 cm cuts from it were made. One was placed in cold HypoThermosol FRS (Sigma-Aldrich, cat no: H4416) at 4°C for a maximum of 24 hours for dissociation and single cell sequencing. The other was snap frozen in an isopentane bath set in dry ice and embedded in OCT medium, while maintaining spatial orientation, and stored at -80°C for cryosectioning and spatial transcriptomics library preparation.

##### 2.2 Myometrium

During delivery of the placenta, the area of the placental bed mirroring the expected placental sampling-site stated above, was established. Three adjacent samples approximately 3mm x 2mm were taken via biopsy, using uterine forceps. Two samples were placed in cold HypoThermosol FRS at 4°C for a maximum of 24 hours, for dissociation and single cell sequencing. The remaining sample was snap frozen in an isopentane bath set in dry ice and embedded in OCT, while maintaining spatial orientation, and stored at -80°C for cryosectioning and spatial transcriptomics.

##### 2.3 Chorioamniotic membranes (CAMs)

From the edge of the placenta closest to the placental tissue sampling site stated above, the membranes were cut from the placental edge. With the chorion facing upward, an approximately 5cm length of membranes was rolled, with the end closest to the placenta being in the middle of the roll. Two adjacent 5cm x 5cm squares of membranes from the sample were placed in cold HypoThermosol FRS at 4°C for a maximum of 24 hours, for dissociation and single cell sequencing. An approximately 1cm length of the roll from the sample was snap frozen in an isopentane bath set in dry ice, embedded in OCT and then stored at -80°C for cryosectioning and spatial transcriptomics.

##### 2.4 Maternal Peripheral Blood Mononuclear cells (PBMC)

Up to 24 hours prior to delivery 7ml of peripheral venous maternal blood was collected in sodium heparin venous blood sampling tube and kept at room temperature for PBMC isolation and single cell sequencing.

#### 3. Metagenomics

Beyond clinical signs of infection (see subsection **1.1 Patient inclusion and exclusion criteria**), we further ruled out infections in our cohort (20 pregnant women, 10 controls and 10 diagnosed with severe early or late PE) with metagenomics. Their PVBP and CAMs samples for each patient were processed across four batches, with the initial batch used as a pilot to optimise the experiments. In addition, and to minimise batch effects, early and late cases and controls were randomised across batches (**Table S1**).

| Patient | Condition | Batch |
|---|---|---|
| **FVB** | Early Control | Pilot |
| **FVQ** | Early Control | |
| **FRK** | Late Control | |
| **FNS** | Late Control | 1 |
| **FEP** | Early PE | |
| **FMM** | Late PE | |
| **FSH** | Late PE | |
| **FJJ** | Early Control | |
| **FYI** | Late Control | 2 |
| **FVT** | Late Control | |
| **FJD** | Late Control | |
| **FAM** | Early PE | |
| **FGS** | Early PE | |
| **FRH** | Late PE | |
| **FLR** | Early Control | 3 |
| **FLJ** | Late Control | |
| **FVS** | Early PE | |
| **FCM** | Early PE | |
| **FFM** | Late PE | |
| **FND** | Late PE | |
| **FCK** | **Positive Control** | **Pilot,1, 2, 3** |

**Table S1**. Batch design for the metagenomics assessment of infections. RNA extraction from Batch 1 for FMM and FJJ was repeated to enhance its quality in Batch 3. FCK is a positive control for infection.

We included an additional patient (FCK) with each batch to assess reproducibility. This sample had clinical signs of a potential infection of unknown origin, which was confirmed in the pilot. It was therefore used as a positive control in subsequent batches. Further, each sample across batches was spiked in with an internal positive control for infection (see below). As a negative control, we used the OCT embedding from the tissue sectioning in a commercial human DNA background, also across batches.

Technical optimizations in the pilot and potential differences arising from the different batches are considered throughout and discussed below.

**3.1 Nucleic Acid Extraction**

Approximately 5 tissue sections each of 5uM thickness for both Placenta Villi and Basal Plate (PVBP) and Chorioamniotic Membranes (CAMs) in each patient were added to DNA/RNA Shield™ and subjected to mechanical disruption. This was followed by parallel DNA and RNA extraction using the ZymoBIOMICS™ DNA/RNA Miniprep Kit. In short, mechanical disruption was carried out using the ZR BashingBead Lysis Tube (0.1 & 0.5 mm), using the following program on an MP Biomedicals™ FastPrep-24 with CoolPrep™ Adapter containing dry ice to reduce sample heat degradation: 1 min at 6.5m/s with 5 min rest, repeated for 3 cycles.

Samples were then briefly centrifuged, removed from the lysis tubes and then spiked with the internal positive control. The pilot batch was spiked with MS2 phage RNA, which is a commonly used internal spike-in for RNA metagenomics. However, in our pilot MS2 was detected in all CAM samples but in only half of the placental samples. This is likely due to the high human RNA content in our placentas, as evidenced by their placental RNA yields, but low preservation, as evidenced by their low RNA Integrity Number (RIN). Such degradation appears to impact the sensitivity for the MS2 phage RNA. In addition, MS2 may also be misclassified as *E. coli* by the `CZID` software tool, which we used to identify and classify pathogens in our metagenomics sequences. Therefore, subsequent batches were spiked with 20uL of ZymoBIOMICS Spike-in Control II (Low Microbial Load) according to manufacturer guidelines.

Importantly, this new spike in contains three bacterial strains not found in the human body, namely: *Truepera radiovictrix*, *Imtechella halotolerans* and *Allobacillus halotolerans*, thus being a better internal positive control and allowing for technical assessment across a range of bacterium types. Further, these bacteria are added as whole cells prior to the bead-beating lysis step, becoming an *in situ* positive control from the beginning of processing and can be used in both DNA and RNA samples.

After this, RNA and DNA extractions were completed following the manufacturer guidelines. These included a DNase treatment for RNA and eluting separately for RNA and DNA. To minimise contamination, samples were always opened and extracted under a class II microbiological safety cabinet. Samples were run in batches of up to 30, and each batch was processed with a known infected CAM sample, acting as an external positive control infected with *Ureaplasma Urealyticum* (and to lesser extent a small amount of reads aligning to *Ureaplasma parvum*). In addition a negative control which contained OCT used for embedding in the cryosectioning process and 40ng/ul of commercial human DNA was used. The controls were used in all steps of processing from bead-beating to sequencing. A Qubit® 4 Fluorometer was used on RNA and DNA for quantification and Genomic DNA and RNA Tapes on the Agilent 4200 TapeStation for integrity assessment.

##### 3.2 Enrichment and library preparation

RNA and DNA libraries were prepared as previously described for Illumina sequencing [3]. Briefly, 500ng of DNA per sample was subject to microbial enrichment using NEBNext® Microbiome DNA Enrichment Kit following the manufactures protocol, except for using double the amount of MBD2-Fc-bound magnetic beads per nanogram (ng) (following recommendation from Great Ormond Street Hospital, GOSH, clinical metagenomics service). The enrichment works by pulling out human CpG-methylated DNA, leaving behind the non-methylated DNA remaining, thus enriched for microbial material. Enriched DNA was quantified as above and was processed with the NEBNext® Ultra™ II FS DNA Library Prep Kit for Illumina. We followed the manufacturer protocol with the following changes: sample input was normalised to 100ng with no size selection, with fragment sizes between 150–350bp.

RNA libraries were prepared using the KAPA RNA HyperPrep Kit with RiboErase according to the manufacturer protocol. This consisted of depletion of rRNA, fragmentation, cDNA synthesis, adapter ligation and amplification. Library inputs were maxed for each sample and PCR cycle number were adjusted depending on input (according to the protocol followed at GOSH clinical metagenomics service). All library preps used IDT xGen™ UDI-UMI Adapters. All steps pre-PCR were carried out under a class II microbiological safety cabinet.

##### 3.3 Sequencing

RNA and DNA libraries from each batch were sequenced either on the Illumina Nextseq 2000 (Pilot and Batch 1) or Illumina Novaseq 6000 (Batches 2 and 3) to achieve between 20M-50M reads per sample. These were demultiplexed and ran through the `CZID` web-based platform for taxonomic profiling [4]. This metagenomics NGS workflow includes data preprocessing and several steps of human read removal by alignment and subsampling. Species were identified by aligning against NCBI databases followed by contig assembly and remapping the contigs. The `CZID` platform outputs a report with the top species hits, including read percentage identity, reads per million and genome coverage. The output from `CZID` was then subject to further filtering using the package `Metathresholds` [3] which compares samples to the negative control with additional thresholds to separate the background from species likely to be truly present.

The degraded quality of most of the PVBP and CAM samples was due to the nature of these tissue types, particularly for the PVBP despite snap freezing within 30 minutes of collection and bead-beating within stabiliser on dry ice. Therefore, sequencing results were interpreted with caution and assessed for presence of pathogenic species on a sample-by-sample basis in addition to the broader thresholds, and immune profiles.

**3.4 Analysis**

After sequencing, reads per sample ranged from 5.7M - 132M. Host (human) read removal and filtering steps in the `CZID` pipeline resulted in a range of 956 - 1.3M reads per sample. Such reduction, above 90%, is typical of low biomass samples[5].

Notably, Batch 1 has lower read-depth and lower Q30 with more reads lost to the low-quality filtering step. However, proportions of both *Ureaplasma* species across batches were similar for CK CAM, our positive control tissue across batches. For this we applied a common measure of dispersion, the Coefficient of Variation as a percentage (CV%), which was below 18.2% for proportion of *Ureaplasma urealyticum* DNA and RNA and *Ureaplasma parvum* DNA but not RNA. This is likely due low abundance, or to the degraded nature of the RNA resulting in misalignment of *U. urealyticum* reads to *U. parvum*. The low CV% suggests that the dominant species *U.urealyticum* is detected in consistent proportion across each run. Therefore it is likely our analyses are not meaningfully impacted by batch effects and can be trusted to identify high load infections across samples regardless of batch.

Further, samples were pooled to equal molarity for sequencing, but there is an inherent heterogeneity across samples due to their differing microbial abundances, which are not known prior to sequencing. These can introduce sequencing biases as higher abundant species monopolise the flow cell cluster generation [6]. Therefore, species reads were normalised for sequencing depth by their reads per million (RPM), which is calculated by `CZID` as the number of sequencing reads aligned to that species in the NCBI database / total reads in the sample and multiplied by 1,000,000. The RPM ratio is then the RPM of the species in a sample divided by the species RPM in the negative control, thus controlling by the species background. Where there are 0 reads for a species in the control, a pseudo number of 1 is used for the division by `CZID`. RPM ratios are commonly used within the field of clinical metagenomics to identify bacterial and viral infections[7].

As a result, one sample was rejected from further analysis (donor FJJ) due to high presence of the bacteria *Ureaplasma parvum* in the CAMs. In RNA: 54,321 reads, 36% of all non-human classified reads in the sample, DNA: 2,957, 32%. When the RNA was repeated at higher read depth in batch 3, this goes up to 871,129 raw reads, 42%. A small number of reads aligned to *Ureaplasma urealyticum* in the CAM DNA sample for FJJ (44, less than 0.05%). Though there were no reads for either of these species in the negative control in batch 1, therefore this could be a misalignment of *U. Parvum*. Our positive control sample was known to contain both species, but cross contamination is not suspected due to zero reads found in any other sample, including the negative control.

As mentioned, our positive control sample, FCK CAM tissues, was included in every batch, with an average fraction of non-human reads, ranging from 40% (DNA) to 42% (RNA), belonging to *Ureaplasma urealyticum*. Thus, our metagenomics approach was sensitive across batches and the samples included in them. Again, some reads were identified for the other *Ureaplasma* species, *U. Parvum (Average 0.1% RNA and 3% DNA)* in the FCK CAM tissues.

Regarding other bacteria, most low abundant ones were known contaminants and skin-related commensals (likely from sample collection). Others could be pathogenic but their low abundance makes their clinical impact uncertain. Although there is no 'standard' RPM-ratio threshold, commonly 10 is used to reduce background [8], which was applied here. In any case, the abundance of any of these bacteria is at least 200-fold below that of pathogenic ones in our positive control, FCK, and 65-fold as the one identified as infected, FJJ. Hence, we consider them background.

Viral infection is more difficult to ascertain using metagenomic sequencing, particularly with low loads. We find low RPM ratios when compared to bacteria, but generally fewer viral reads are needed to indicate an infection. The number of viral reads per sample range across all batches from 0 to 75 reads, with control reads of each species ranging from 0 to 69. Whereas infected samples with bacteria have tens of thousands of bacterial reads, with even background bacteria commonly presenting hundreds, or even a few thousand of sequencing reads.

Still, we find sample FEP to have HHV-6 and HHV-7 viruses in its DNA and RNA, suggesting not only a latent infection, as is typical for HHV-6, but also potential active expression of the virus. This is, however, at low level as we found only two RNA reads in both FEP CAMs and PVBP (RPM of 0.038 and 0.048 respectively), agreeing with the lack of clinical symptoms and an extremely low-level active infection if present. There have been recent studies investigating a potential association between HHV-6 infection and risk of preeclampsia [8, 9] , although its significance differs between these studies. It should be noted that after processing, HHV-6 and 7 was found to be present in the commercial donor DNA used as a background for the negative control in each batch (between 0-8 reads across batches). Presence was confirmed in DNA samples by qPCR by the GOSH clinical metagenomics service (not shown).

Similarly, we found sequencing reads aligning to Covid-19 in many of the samples (FSH, FVB, FMM, FVQ, FAM, FCM, FGS, FJD, FLR, FND, FPYI, FRH, FVS, FVT, FJJ), although very few reads and low RPM ratios per sample. For batches 1 and 2 (for samples FSH, FMM, FJJ, FVB, FVQ), the number of reads ranged from 1 to 4, with 0 to 1 read in the negative control. For the final batch 3 (for samples FAM, FCM, FGS, FJD, FLR, FND, FYI, FRH, FVS, FVT), the number of reads was slightly higher, ranging from 1 to 17 but the negative control had a range of reads from 2 to 69. We also find reads aligning to Cytomegalovirus (CMV) (for samples FAM, FCM, FGS, FJD, FJJ, FLJ, FLR, FND, FYI), ranging in number from 1 to 5, with negative control reads ranging from 0 to 2. In addition we find reads aligning to Pegivirus hominis in sample FGS (21 reads in CAM RNA, 10 in PVBP RNA) traditionally thought to be non-pathogenic but gaining attention in pregnancy studies as it has been found in placenta and decidua. However, we conclude that there is uncertain clinical significance from these virus reads alone, therefore decision on these samples should be based on immune profiling and clinical metadata. From this combination, we conclude there is not enough evidence for infection.

Finally, we generated a PCoA (Principal Coordinate Analysis) plot based on the Bray-Curtis dissimilarity for for the combined two tissues per sample. This allows us to visualize the relationships between samples from their taxonomic composition, specifically using the non-human proportions for each sample. Importantly, control and severe PE samples cluster together, separated by their different spike-ins (different between the pilot and the subsequent three processing batches). Samples falling outside both these clusters, CK (positive control) and FJJ, are infected with *Ureaplasma*. Thus, aside from sample FJJ that was subsequently excluded, infection is deemed likely not present to a level that confounds our analyses of the molecular signatures of severe PE, including immune ones, with cases and controls being generally undistinguishable from their metagenomics profiles.

#### 4. Single cell and spatial transcriptomics experiments

##### 4.1 Single cell tissue dissociation

Cell dissociation and death cell removal prior library preparation for each sampled tissue type are detailed below.

**PVBP and CAM**. Samples were removed from HypoThermosol FRS and transferred to C-Tubes (Milteny Biotec) on ice before being minced with scissors. 10ml of Accutase (Sigma-Aldrich) was then added to the C-Tubes. The tissues were then dissociated on a gentleMACS Octo dissociator using a custom protocol:

1. Loop 6 x
2. Spin 200 rpm, 4”
3. Spin -200 rpm, 2”
4. Spin -20 rpm, 2’ 30”
5. End loop
6. End

Dissociated cell suspensions were then passed through stacked 100um and 70um cell strainers (Milteny Biotec), pre-wetted with 2ml of DMEM, 10% FBS. Cell strainers were then rinsed with a further 10ml of DMEM, 10% FBS. The two samples of CAM tissue were combined into one suspension.

Cells were pelleted by centrifugation at 300RCF for 5 minutes. The supernatant was then removed, and the pellets were resuspended in 20ml of Red Cell Lysis buffer (Milteny Biotec) and incubated for 10 minutes at room temperature on a rocker. Then 30ml of cold 1x DPBS was added, and cells were pelleted by centrifugation at 300RCF for 5 minutes. The supernatant was removed, and cells were washed by resuspending in 5ml DMEM, 10% FBS before being pelleted by centrifugation at 300RCF for 5 minutes.

The supernatant was then removed, and cell pellets were resuspended in 1ml of Dead Cell Removal MicroBeads and incubated for 15 minutes at room temperature. Suspensions were then loaded onto MACS LS columns (Miltenyi Biotec) pre-rinsed with 3ml 1x Binding Buffer. Columns were rinsed with a further 3ml of Dead Cell Removal 1x Binding Buffer. Cells were then pelleted by centrifugation at 300RCF for 5 minutes. The supernatant was removed, and cells were resuspended in 1ml DMEM, 10% FBS.

In addition, nuclei single cell suspensions were prepared for the CAM from donors “FVQ”, “FCM”, “FGC”, “FLJ”, “FRK”. Briefly, snap frozen and OCT embedded CAM samples were cryosectioned on a Bright OTF5000 cryostat. Ten 25um sections of tissue were collected.

Nuclei were isolated from the sections using Chromium Nuclei Isolation Kit (10X Genomics), following 10X Genomics user guide CG000505.

**Myometrium**. The two myometrium samples were removed from HypoThermosol FRS and transferred into one C-Tube (Milteny Biotec) on ice before being minced with scissors. 5ml of Accumax (Sigma-Aldrich) was then added to the C-Tube. The tissues were then dissociated on a gentleMACS Octo dissociator using the same PVBP and CAM custom protocol (see above). Dissociated cell suspension was then passed through stacked 100um and 70um cell strainers (Milteny Biotec), pre-wetted with 2ml of DMEM, 10% FBS. Cell strainers were then rinsed with a further 5ml of DMEM, 10% FBS.

Cells were pelleted by centrifugation at 300RCF for 5 minutes. The supernatant was then removed, and the pellet was resuspended in 5ml of Red Cell Lysis buffer (Milteny Biotec) and incubated for 10 minutes at room temperature on a rocker. Then 10ml of cold 1x DPBS was added, and cells were pelleted by centrifugation at 300RCF for 5 minutes. The supernatant was removed, and cells were washed by resuspending in 1ml DMEM, 10% FBS before being pelleted by centrifugation at 300RCF for 5 minutes. The supernatant was removed, and the cell pellet was resuspended in 250ul of 1x DPBS, 0.04% BSA and passed through a 40um Flowmi Cell Strainer (Sigma-Aldrich).

**PBMCs**. Seven ml of peripheral venous maternal blood was carefully layered over 3ml of Histopaque 1077 (Sigma-Aldrich) and centrifuged at 400RCF for 30 minutes with the centrifuge acceleration and brake at their lowest settings. The phase containing the plasma was removed. The layer of PBMCs was removed and the cells were washed with 5ml of RPMI and then pelleted by centrifugation at 300RCF for 5 minutes. The supernatant was removed, and the cells were resuspended in 1ml RPMI.

##### 4.2 Single cell library preparation

Dissociated cells, isolated PBMCs and isolated nuclei were checked for concentration and viability using an Acridine Orange/Propidium Iodide Stain (logos Biosystems) on a logos Biosystems Luna FL automated cell counter.

Single cell suspensions per tissue and donor were used to generate single cell/nuclei transcriptomic and single cell/nuclei TCR libraries. We loaded 20,000 cells/nuclei into a 10X Genomics Chromium Controller using the Chromium Next GEM Chip K and Chromium Next GEM Single Cell 5' Kit v2 kits (10X Genomics) as per 10X Genomics user guide CG000331.

##### 4.3 Spatial transcriptomics library preparation

Ten-micrometre sections of snap frozen PVBP, Myometrium and CAM samples were loaded onto Visium Spatial Gene Expression slides (10X Genomics) as per 10X Genomics protocol

CG000240. The area of the PVBP samples chosen for loading onto the relevant capture areas contained the edge proximal to the decidua basalis. Loaded sections on Visium slides were subsequently methanol fixed, stained with Haematoxylin and Eosin (H&E) and imaged as per 10X Genomics protocol: GC000160 on a Motic EasyScan One slide scanner, at 40X magnification. Following 10X Genomics user guide CG000239, sections were then subjected to permeabilization for mRNA capture, reverse transcription and second strand synthesis to generate full length cDNA, followed by library preparation. Permeabilization times of 18 minutes for CAM and Myometrium and 6 minutes for PVBP, were established as appropriate using the Visium Spatial Tissue Optimisation kit (10X Genomics) following 10X Genomics protocols and user guide CG000240 for sectioning and slide loading. We followed manufacturers protocol GC000160 for H&E staining and brightfield imaging and protocol CG000238 for mRNA capture, reverse transcription, second strand synthesis and fluorescent imaging.

A tissue section per patient was loaded in one of the four Visium capture areas.

**4.4 Sequencing**

Resulting spatial transcriptomic, single cell and single nuclei transcriptomic and T-Cell Receptor libraries were sequenced on an Illumina Novaseq 6000 using five S4 (200 cycle) v1.5 sequencing kits (Illumina), with a configuration of Read 1: 28 cycles, Index read 1: 10 cycles, Index read 2: 10 cycles, Read 2: 190 cycles. Libraries were sequenced to a minimum coverage of 20,000 reads per cell or 25,000 reads per Visium spot.

### 5. Immunofluorescence assay

#### 5.1 Sectioning and fixation

Early PE and Early Control PVBP samples were cryosectioned on a Leica CM1950, generating 8μm sections. All PVBP sections were loaded onto one glass slide. A separate PVBP sample was loaded onto a second slide to be used as a negative antibody control. Slides were incubated at 37C for 1 minute before fixation in 10% Formaldehyde for 30 minutes at room temperature. Slides were washed in 1X DPBS for 1 minute followed by 1% SDS for 2 minutes and a further two washes in 1X DPBS for 1 minute each. Slides were then immersed in 70% methanol for 60 minutes on ice before being washed twice in 1X DPBS for 1 minute each.

#### 5.2. Blocking, permeabilization and staining

Each slide was then washed with 500ul 1X DPBS 0.05% Tween-20, before adding 500ul blocking and permeabilization buffer consisting of: 1X DPBS 0.1% Tween 10%FBS 0.1% Triton-X with 10mg/ml dextran sulphate and incubating at room temperature for 60 minutes. Staining was performed by adding 500ul staining buffer consisting of 1X DPBS 0.1% Tween 10%FBS with 10mg/ml dextran sulphate and a 1:100 dilution of CoraLite® Plus 488-conjugated MX1 Recombinant antibody (Proteintech). Staining buffer without antibody was used for negative antibody control. Slides were incubated overnight at 4C in the dark. Following staining, slides were washed three times with 500ul 1X DPBS 0.05% Tween-20 for 10 minutes at room temperature each. Then, 500ul DAPI at 5ug/ml was added for 1 minute before being washed three times with 500ul 1XDPBS 0.05% Tween-20 for 1 minute each. A coverslip was then added using SlowFade Gold Antifade mounting medium.

#### 5.3 Imaging and positive cell quantification

Stained slides were imaged on a Nikon Ti2 inverted microscope at 20X magnification. Resulting image files were loaded into `QuPath 0.5.0`. Fluorescence thresholds for exclusion of background and autofluorescence were established using the negative antibody control. A morphological region was delineated to include an approximately equivalent proportions of decidua and villous placenta to capture both maternal and fetal contributions. Sections for FGS, FVQ and FVS lacked the decidual regions and were therefore excluded from further analysis. Positive cell detection analysis was performed using DAPI for nuclei detection with a 5μm cell expansion and a nuclear, maximum intensity threshold for detection of MX1 positive cells. `RStudio` was then used to generate comparative plots.

#### 6. Quantification of IFN-I expression in PBMCs

PBMCs isolated from whole blood, as described above, were subjected to RNA extraction using the Qiagen RNEasy plus kit (see **Extended Data Table 2** for donor and sample details). RNA extracts were quantified by Nanodrop and normalised to a concentration of 2ng/ul. Extracts where then subjected to qPCR using taqman probes for 4 interferon related genes (IFIT1, IFI44L, ISG15, RSAD2) along with 1 housekeeping gene (18S). The NEB Luna® Universal Probe One-Step RT-qPCR Kit was used with an input of 1ul of sample added to 4ul of resulting qPCR mastermix. qPCR reactions were performed using an Applied Biosystems Quantstudio 5 Real-Time PCR instrument, using cycling conditions as per recommended for the reaction kit. Each sample was run in triplicate along with 1 negative template control on each reaction plate. Technical replicates that deviated from the remaining two by over 1 Ct (Cycle threshold; number of cycles necessary to replicate enough DNA/RNA for detection) were excluded as outliers. Samples where appropriately randomised to ensure the presence of both cases and control on each reaction plate.

Output files where initially processed using the Design & Analysis Software by Applied Biosystems with Ct values per replicate group exported as a .csv file. Replicate group Ct .csv files were then compiled for all samples using `Microsoft Excel`. To normalise gene expression levels across samples DCt (Delta CT; difference in cycle threshold between a target and reference gene) values were calculated per sample for each of the four interferon related genes against the housekeeping gene. The mean DCt value of the control samples was calculated for each gene and the DDCt (Delta Delta CT; DCT difference between target and reference samples) values per sample for each gene were then calculated against the mean DCt control value to obtain relative gene expression values. DDCt values were then converted into fold change values as per 2^DDCT. Mean and median values of the four interferon related genes per sample were calculated. Wilcoxon tests were applied in `RStudio`, comparing cases and controls for each gene, along with the average and median interferon gene fold change.

#### 7. Single-cell transcriptomic analysis

##### 7.1 Pre-processing: alignment, count matrix and ambient RNA correction

For each single cell and single nucleus library, the raw bcl files were converted into fastq files using `mkfastq` (version `cellranger-v7.1.0`; 10X Genomics). Sequencing reads were aligned to the GRCh38-2020-A human reference genome (GENCODE v32/Ensembl98 distributed by 10X Genomics) and a matrix of unique molecular identifier (UMI) counts and TCR clonotypes per library was obtained using `cellranger` "multi". We exported the first 10,000 cells ranked as true cells (using the --forcecells option). We used `CellBender` v0.2.0 to correct for ambient RNA per library and classify cell-containing droplets from empty ones [10].

##### 7.2 Fetal and maternal single cell classification

For cell classification into fetal or maternal origin, we used *`Freemuxlet`* v.1 as implemented in the `popscle` software (https://github.com/statgen/popscle), designed to demultiplex and genotype single-cell RNA sequencing data from mixed samples [11]. To obtain the genotype per cell, we first acquired a variant site list (in VCF format) with the most common SNPs from the 1000 Genomes Project, serving as the panel of SNP positions. Then, we used the *dsc-pileup* tool as implemented in `popscle` to identify the allelic composition, base quality, and the number of reads from each allele at each common SNP location per library using the BAM file output from the 10X Genomics `cellranger` "multi" pipeline described above. We also used the option *--group-list* to run `dsc-pileup` focusing on the droplets that most likely represent true cells from `CellBender` output.

Before running `Freemuxlet` classification analysis, we first combined the `dsc-pileup` outputs from the tissue PVBP and either the Myometrium, PBMCs or CAM per patient. This computational mixing assures that both fetal/maternal fractions are well represented in each classification analysis (circumventing the low abundance or absence of fetal cells in the Myometrium and PBMC).

Then, we ran `Freemuxlet` with default values and two groups (*i.e* two individuals: *--nsample 2*). Based on the SNP profiles, this analysis assigns each cell a genotype: 0, 1 or 0/1 (i.e doublets containing reads from two individuals). To identify the inferred genotype that corresponded to the fetus or mother, we calculated the average expression of a set of trophoblast markers ("CGA", "CYP19A1", "GH2", "PAPPA", "VGLL1", "PAPPA2", "HLA-G") per genotype. We assigned as fetal the genotype with higher average trophoblasts expression. Additionally, we verified the `Freemuxlet` genotype assignment by correlating the SNPs obtained using a DNA microarray (Infinium Global Diversity Array v1.0) on a subset of samples.

We genotyped 8 pairs of fetus and maternal donors using DNA isolated from cord blood and PBMC respectively. These were positively correlated with the SNPs genotypes inferred by `Freemuxlet`, corroborating the assignment of fetal origin by expression of trophoblasts markers.

##### 7.3 Single cell Quality Control (QC)

As part of our QC, doublets were detected in two ways: 1) using `Scrublet` v0.2.3 [12] (*manual threshold = 0.25*) per library; and 2) as a droplet with a mixed genotype (0/1). These were filtered out.

QC metrics, including the number of detected genes, the number of unique molecular identifiers per cell, the percentage of mitochondrial genes and percentage of haemoglobin genes per cell were calculated using `Seurat` v4.2.1 [13]. Thresholds were selected after inspecting the distribution of these metrics. Cells were filtered out if: 1) number of detected genes was below 400 and above 6,000; 2) number of unique molecular identifiers was above 30,000; and 3) percentage of mitochondrial reads was more than 12% and percentage of haemoglobin genes was more than 0.25.

##### 7.4 Single cell integration and cell type annotation

After pre-processing, gene expression matrices from all donors and all tissues were integrated using `Seurat` v4.2.1 [13]. Briefly, a standard workflow including normalisation ("LogNormalise"), scale ("ScaleData") and highly variable genes selection (2000 genes) were used to perform principal component analysis (*npcs = 25*), k-nearest-neighbour calculation (*k.param = 20*) and graph-based community detection using Louvain clustering. `Harmony` v0.1.0 [14] was used to correct for batch effect due to 10X method used (single cell or single nuclei).

We annotated cell types using gene expression of manually curated genes from the literature and genes differentially expressed per cluster (obtained using `Seurat` function "FindMarkers"). Cell types were annotated at three levels of resolution (refereed as annotation level 1, level 2 and level 3). First, a coarse cell group annotation rendered six major groups: 1) Trophoblasts; 2) Stromal; 3) Vascular; 4) Myeloid; 5) B; and 6) T/NK cells at annotation level 1. Subsetting each of these groups and repeating the clustering pipeline described above allowed the assigment of two more levels of annotation with increasing granularity. During clustering, we detected further doublets (identified by a mixed profile of gene expression in the cluster) and those were also filtered out.

In all the sections below, cell types from donor FJJ were excluded as explained in section *1.1 Patient exclusion and inclusion criteria*.

##### 7.5 Cell abundance differences from disease and gestational age

To measure the differential abundance of cell populations at single-cell resolution due to disease or gestational age, we used `dawnn` v1.0.8 [15]. Briefly, we computed the probability of each cell to be differentially abundant in each neighbourhood using a pretrained deep neural network model [15].

Using as an input a `Seurat` object per tissue, we calculated a KNN graph using the first 10 principal components (*reduced_dim* = *pca*) for the PVBP, Myometrium and PBMCs tissues. For the tissue CAM, we used `Harmony` v0.1.0 [14] (*reduced_dim* = *harmony)* to correct for batch effects introduced by the two different single cell methods (single cell gene expression and single nuclei gene expression) and used the first 10 components from harmony. We applied a false discovery rate correction with the Benjamini-Yekutieli procedure as implemented in `dawnn` v1.0.8 using an *alpha = 0.05*.

We tested for differentially abundance separately comparing: 1) conditions (Control or PE); or 2) gestational age (Early or Late). We measured whether a cell is called differentially abundant in one comparison, in both or in neither.

##### 7.6 Cell type differential expression analysis per tissue

We tested for single cell differential expression in each tissue while accounting for gestational effects and donor-to-donor variability. Cell types with fewer than 25 cells per condition or present in only one donor were excluded from the differential expression analysis. Genes lowly expressed (fewer than 20 total counts) were also filtered out.

For each tissue, we first aggregated the measured counts per donor and cell type at annotation level 3 into a pseudobulk expression profile (as recommended in [16]). Each pseudobulk belonged to a unique condition *c,* specified as:

$$c \in \{Early_{control},\ Early_{PE},\ Late_{control},\ Late_{PE}\}.$$

We modelled the cell-type specific pseudobulk mRNA counts $y_{gd}$ of gene $g$ from donor $d$ as a negative-binomial generalised linear model (NB-GLM):

$$y_{gd} \sim NB(\mu_{gd},\ \phi_{gd}), \qquad \text{(eq.1)}$$

where $\mu_{gd}$ is the mean number of counts of gene $g$ in donor $d$ and $\phi_{gd}$ is the dispersion parameter estimated using conditional likelihood implemented in `EdgeR` v3.36.0 [17].

The effect of gestational age and disease on the counts of each gene in each condition can be modelled using a log-linear model [18]. Thus, the expected log count value $\mu_{gd}$ is given by:

$$\log\mu_{gd} = a_1\beta_g^1 + a_2\beta_g^2 + a_3\beta_g^3 + a_4\beta_g^4 + \log N_d \quad \text{(eq.2)}$$

where $a_i$ is an indicator variable for condition: $a_1$ =1 for $Early_{control}$, $a_2$ =1 for $Early_{PE}$, $a_3$ =1 for $Late_{control}$, $a_4$ =1 for $Late_{PE}$, and equals zero otherwise. Following the same numbering scheme, $\beta_g^i$ is the regression coefficient associated with gene $g$ for condition $i$, e.g. $\beta_g^1$ is the regression coefficient associated with gene $g$ for $Early_{control}$.

$N_d$ is the total number of counts from donor $d$.

Using `EdgeR`, we used a full rank matrix with donors as rows and conditions (called the variable $c$ ) as columns to fit the above log-linear model (eq.2) using the design matrix: $\sim 0 + c$ .

Modelling gene expression using this parameterisation allows us to separate the effects of gestational age and disease and test for differences in gene expression per gene $g$ by using a linear combination of the regression coefficients [18] (via a contrast matrix) such as:

(1) Early disease effect is given by:

$$EPE_g = \beta_g^2 - \beta_g^1 \quad \text{(eq.3)}$$

corresponding to the contrast $Early_{PE}$- $Early_{Control}$

(2) Late disease is given by:

$$LPE_g = \beta_g^4 - \beta_g^3 \quad \text{(eq.4)}$$

corresponding to the contrast $Late_{PE}$ - $Late_{control}$

(3) The average PE effect is given by:

$$APE_g = \frac{\beta_g^2 + \beta_g^4}{2} - \frac{\beta_g^1 + \beta_g^3}{2} \quad \text{(eq.5)}$$

corresponding to the contrast: ($Early_{PE}$ + $Late_{PE}$)/2 - ($Early_{Control}$+ $Late_{control}$)/2

(4) Finally, the average gestational effect is given by:

$$AGA_g = \frac{\beta_g^1 + \beta_g^2}{2} - \frac{\beta_g^3 + \beta_g^4}{2} \qquad \text{(eq.6)}$$

corresponding to the contrast: $(Early_{PE}$ + $Early_{Control})$/2 - $(Late_{PE}$ + $Late_{control})$/2

We tested for differential expression per contrast using a likelihood ratio test (LRT) implemen ted in `EdgeR` v3.36.0 [17] (as recommended by [16]). Multiple testing was controlled by using

the Benjamini-Hochberg (BH) correction. Genes were considered differentially expressed if the log fold-change > 0.5 and the FDR < 0.05.

**7.7 Macrophages differential expression across tissues**

We also tested for differences in gene expression of Macrophages in severe preeclampsia using a GLM framework (`EdgeR`). Each donor has samples from all tissues, thus we modelled gene expression using a paired design to adjust for donor-specific effects using the design matrix: $\sim 0 + tissue + donor$. We selected only Macrophages of Early PE or Late PE to compare macrophages across tissues only in disease.

As before, we used a log-linear model for tissues $k \in \{PVBP, Myometrium, CAM\}$:

$$\log \mu_{gd} = t_k \beta_g^k + h_d \gamma_g^d + \log N_d \quad \text{(eq.7)}$$

Where:

$t_k$ is an indicator variable equal to 1, if sample is from tissue *k,* 0 otherwise;

$\beta_g^k$ is the regression coefficient associated with tissue-specific log-expression (as in eq. 2);

$h_d$ is an indicator variable equal to 1, is sample is from donor $d$, 0 otherwise;

$\gamma_g^d$ captures to donor-specific effects;

$N_d$ is the total number of counts from donor $d$.

We compared gene expression between PVBP and Myometrium and PVBP and CAM using the linear combinations: 1) $\beta_g^{PVBP} - \beta_g^{Myometrium}$ and 2) $\beta_g^{PVBP} - \beta_g^{CAM}$, respectively. Finally, we used a likelihood ratio test (LRT) and a Benjamini-Hochberg (BH) correction as before.

**7.8 Cell proportions per tissue**

To test for differences in cell proportions accounting for donor variability and gestational age, we used the GLM framework implemented in the *propeller* function of the R package `speckle` v0.0.3 [19]. Briefly, the proportion of cell type $j$ in a specific tissue from donor $d$, is given by:

$$p_{jd} = \frac{x_{jd}}{N_d} \text{ , (eq.8)}$$

where $x_{jd}$ denotes the number of cells observed for cell type $j$ in a specific tissue from donor $d$ and $N_d$ is the total number of cells sampled in a specific tissue per donor. We perform a variance stabilising step by applying an arcsin square root transformation $z_{jd}$ implemented in *propeller* [19] as:

$$z_{jd} = \arcsin(\sqrt{p_{jd}}) \quad \text{(eq.9)}$$

For tissue $k \in \{PVBP, Myometrium, PBMC\}$, the mean expected transformed proportion per cell type $z_{jd}$ can be modelled as:

$$E(z_{jd}) = a_1\beta_j^1 + a_2\beta_j^2 + a_3\beta_j^3 + a_4\beta_j^4, \text{ (eq.10)}$$

similarly to eq.2, $a_i$ is an indicator variable for condition and $\beta_j^i$ is the regression coefficient associated with the transformed cell proportions per cell type $j$. As above, we use a matrix with donors $d$ as rows and conditions $c$ as columns to fit the above linear model using the design matrix: $\sim 0 + c$ .

For tissue $k = CAM$, we added a term to correct for the effect introduced by using two different library preparation methods (single-cell gene expression and single-nuclei gene expression) as:

$$E(z_{jd}) = a_1\beta_j^1 + a_2\beta_j^2 + a_3\beta_j^3 + a_4\beta_j^4 + f_1\beta_j^5 \text{ (eq.11)}$$

Here $f_1$ is the variable corresponding to a given method (0 = single-cell or 1 = single-nuclei) associated to donor $d$ and $\beta_j^5$ is the regression coefficient encoding the effect of library preparation protocol. This model uses the modified design matrix: $\sim 0 + c + method$.

After model fitting, we tested for differences in cell proportions per cell type using a moderate t statistical test for each comparison as implemented in `propeller` [19] . As before:

(1) Early disease is given by:

$$EPE_j = \beta_j^2 - \beta_j^1 \qquad \text{(eq.12)}$$

corresponding to the contrast $Early_{PE}$- $Early_{Control}$

(2) Late disease is given by:

$$LPE_j = \beta_j^4 \text{ -}\beta_j^3 \quad \text{(eq.13)}$$

corresponding to the contrast $Late_{PE}$ - $Late_{control}$

We applied a false discovery rate correction using BH procedure. After statistical testing, only cell types that came from one donor or only present in one condition were disregarded.

**7.9 Differential cell abundance and gene expression in pseudotime**

To identify cell populations associated with severe preeclampsia and their transcriptional profiles, we quantified the differential abundance of cell types in cases compared to controls across their pseudotime trajectories. We focused on syncytiotrophoblast (STB) cell populations expressing hypoxia-related genes (HTRA1, HTRA4, FSTL3, ENG, TMEM454), macrophages expressing inflammation-related genes (IL10, GSDMA). This pseudotime analysis can identify continuous changes of abundance in a given cell type that correlate with changes in gene expression.

Cell type populations were analysed by fitting them into a pseudotime trajectory, with differences in cell abundance between severe PE and control cases and the mean gene expression per cell quantified at equal pseudotime intervals. Briefly, a UMAP was generated for the STB cells (including progenitors cytotrophoblasts, CTB) and a UMAP for macrophages (function as implemented in Seurat v.4.3.0). We used `slingshot v.2.6.0` [20] to infer cell trajectories in pseudotime following the `Condiments` pipeline [21]. For the STB, we used the annotation level 3 and set as starting point the CTB proliferating cluster, while no initial cluster was set for macrophages. For visualisation, we calculated abundance imbalance scores (control vs severe PE) across the UMAPs using the *imbalance_score* function implemented in `Condiments`.

To identify pseudotime intervals containing differential abundance of cells of any condition (control or severe PE), considering sample variation, we used in-house scripts to generate a sequence of pseudotime intervals of *n* pseudotime units (we divided the maximum pseudotime by 10 to obtain *n*). For each interval, we obtained the number of cells from each donor (from the sequencing library) or the proportion of cells relative to the donor's total number of cells for a cell type. We then performed a Wilcoxon rank-sum test to determine if the distribution of cell counts or proportions of the two groups of samples (conditions) is different, and applied Benjamini-Hochberg correction for multiple testing (threshold 0.2). This test was performed only at intervals where cells from three or more donors per condition were present, separately for early and late disease. Significant results were only possible when many donors from the same condition had the highest cell counts

To test whether hypoxia-related genes are upregulated in STB populations that are more abundant in PE, we calculated the mean gene expression per cell for hypoxia-related genes (defined as the sum of the expression of all such genes) at each pseudotime interval. We then correlated the change in gene expression along pseudotime with the difference in cell abundance (*corr.test* function as implemented in R), where cell abundance was defined as the difference between the mean number of cells per donor in severe PE cases and controls within each interval. Similarly, to identify genes up- or down-regulated in macrophages populations that are more abundant in PE compared to the rest of macrophages, we correlated difference in cell abundance along pseudotime with mean gene expression per cell at each pseudotime interval for each of the immune-response related genes VEGFB, MERTK, SPP1, CXCR4, IL10, MSR1, FN1, CTSA, CTSD, TYMP, GSDMA, VCAN, RMDN3, TGFB, MRC1, FOLR2, C1QB, CSF2, PDGFD, CD163, LYVE, and TREM2 separately.

##### 7.10 Cell-to-cell communication

`MultiNicheNet` (multinichenetr v1.0.3) [22] was used for differential ligand-receptor analysis in Early and Late disease. Cell types from the tissues PVBP and Myometrium were integrated into a single `Seurat` object to take advantage of their spatial continuity. Cell types at annotation level 3 with less than 25 cells and coming from one single donor were excluded. `Nichenet` v2 ligand-receptor network and matrices from "*organism* = *human*" were obtained from Zenodo (DOI: 10.5281/zenodo.7074291).

Briefly, `MultiNicheNet` runs a differential expression analysis per cell type followed by a prioritisation score of ligand-receptor pairs accounting for donor expression levels and ligand intracellular activity [22]. Early disease and Late disease were modelled using the same GLM as before (eq.3 and eq.4). We applied a p-value cutoff of 0.05 and log-fold change threshold of 0.5. Other parameters were set as default. We selected the top 150 targets per ligand and visualise the top predictions using `MultiNicheNet` inbuilt functions.

#### 8. Spatial transcriptomics analysis

##### 8.1 Alignment and gene counts per spot

For the Visium spatial transcriptomics, demultiplexing, alignment of reads to the human reference genome (GRCh38-2020-A, GENCODE v32/Ensembl98 distributed by 10X Genomics), identification of spots for each tissue and quantification of gene counts per spot were performed using `spaceranger` v2.0.0, as distributed by 10X Genomics.

##### 8.2 Spatial domain identification per tissue

Spatial domains for the Visium spatial transcriptomic results were independently identified by two researchers using the 10X Genomics `Loupe Browser` 7. Spatial domain annotations were cross referenced to generate a consensus annotation.

First, we used a *kmeans* clustering of the gene expression per capture area (k=4 for CAM and k=3 for PVBP and Myometrium tissues) to generate clusters which broadly aligned with the morphologically expected tissue domains. We ran a differential expression analysis (in-built in Loupe browser) to compare gene expression between clusters and inspected the presence of cell type markers with known spatial distribution (i.e. decidual cells in the decidua basalis and decidua parietalis from PVBP and CAM, respectively). We then used this information and prior knowledge of tissue morphology to manually annotate spatial domains.

We also manually identified Visium spots with poor quality or low-confidence spatial domain assignment and labelled them as “background”. Briefly, a spot was classified as “background” when it belonged to the following categories:

- Tissue processing artefacts:
    - Tissue folds.
    - Areas of tissue damaged during snap freezing.
- Technical artefacts:
    - Areas spanning multiple spatial domains with exceptionally low UMI capture per spot.
    - Spots that were assigned as under tissue by `spaceranger` v2.0.0 in error due to staining residue on slide.
- Ambivalent annotation:
    - Areas that could not be confidently assigned to a particular spatial domain using the methodology outlined above.

Visium spots belonging to “background” were excluded from further analysis.

##### 8.3 Spatial hypoxia gene expression score in placenta villi

We calculated an “hypoxia score” using the `Seurat` function *AddModuleScore* [13]. Briefly, this function calculates the average expression levels of a set of genes per spot subtracted by the aggregated expression of control genes. The module score was calculated on spots belonging to the spatial domain “villi” per capture area. The hypoxia score included the genes: "HTRA1", "HTRA4", "FSTL3", "EGLN3", "TMEM454” and we used 30 control genes.

##### 8.4 Cell type deconvolution

We used `cell2location` v0.1.3 [23] to infer cell type abundances per spot in Visium spatial transcriptomics. We trained a single cell regression model using the cell type annotation at level 2 as a single cell reference (excluding donor FJJ), keeping genes expressed in at least 3% of the cells and excluding genes expressed in less than 5 cells. We trained the single cell regression model for 350 epochs.

To deconvolute the cell types per spot, we set the parament *N_cells_per_location = 8* and a *detection_alpha = 20*. We trained the deconvolution model for 3,000 epochs and exported the 5% quantile of posterior distribution for visualisation as recommended in `cell2location` documentation. Each Visium capture area was deconvoluted separately.

##### 8.5 Spatial expression statistics using a mixed-effects model

To test for differences in abundance of EVT per spot in Early or Late disease, we used a linear mixed-effects model implemented in the packages `lme4` v1.1-27.1 and `lmerTest` v3.1-3. Briefly, we considered the condition (Early Control vs Early PE or Late Control vs Late PE) as the fixed effect while the variation between donors belonging to the same condition was considered a random effect using the model:

$$lmer((c2loc\_EVT) \sim condition + (1 \mid donor)) \quad \text{(eq. 13)}$$

##### 8.6 Spatial correlation of EVT and SC-CTB in CAM

We used a Pearson correlation to quantify spatial distribution of EVT and SC-CTB in the CAM spatial domain “chorion”. We filtered out spots with SC-CTB abundance less than 0.5. For statistical comparison between conditions in Early or Late disease, we used a Fisher’s Z-transformation and computed a p-value (two-tailed test).

##### 8.7 Neighbourhood analysis in the CAM

To quantify the spatial arrangement of spots with high abundance of the cell type SC-CTB (abundance greater than 3) in the spatial domain “chorion” (CAM), we calculated the identity of their neighbour spots using the nearest neighbour algorithm (*k=2*) as implemented in the `RANN` v2.6.1 package.
